# Minimal, synthetic arginine deiminase pathway provides full ATP demand for cellular growth

**DOI:** 10.64898/2026.11.14.766618

**Authors:** Karin Schann, Joep Houkes, Gea K. Schuurman-Wolters, Sara Benito-Vaquerizo, Tessa de Noojier, Yining Li, Emma van Aalzum, John van der Oost, Bert Poolman, Nico J. Claassens

## Abstract

Bottom-up construction of synthetic cells requires a simple, yet robust ATP-generating system. The arginine deiminase (ADI) pathway offers a minimal, enzyme-based alternative to more complex ATP-producing systems such as glycolytic routes, respiratory chains or photosystems. Here, we implement the ADI pathway in *Escherichia coli* and test its capacity to fully replace native ATP generation. First, we designed and tested synthetic ADI operons through combinatorial library screening and selected high-performing variants. Using the best performing-ADI variant, we progressively reduced the cells’ dependence on native ATP generation while increasing selection for ADI-dependent growth through targeted knockout strains and growth conditions, followed by adaptive laboratory evolution. This resulted in a strain in which the ADI pathway sustained growth in minimal medium at a doubling time of 21 hours, serving as the sole source of ATP. Whole-genome sequencing of evolved strains revealed that this transition required loss-of-function mutations in the native respiratory chain. These findings demonstrate that the ATP supply of a cell can be fully supported by the ADI pathway, which has not been demonstrated in nature. Moreover, the ability of the pathway to sustain growth with a relatively high growth rate highlights its potential to fully power synthetic cells and support other biotechnological applications.

## 1. Introduction

The generation of metabolic energy is among the most fundamental and conserved processes in living systems. At the cellular level energy is stored in two interconvertible forms: in high-energy phosphate bonds, mostly ATP, and as electrochemical ion gradients across membranes, which are used to drive endergonic reactions and processes [1, 2]. The classical example of metabolic energy conservation is oxidative phosphorylation in which the free energy of the oxidation of e.g. NADH is used to pump protons across the membrane, and the formed electrochemical proton gradient (or proton motive force) is used to drive ATP synthesis via F_0_F_1_-ATP synthase [3]. Although highly efficient, respiration is among the most complex processes in living cells and involves many proteins and cofactors. By contrast, ATP can also be generated by substrate-level phosphorylation reactions, which do not require a membrane apparatus and can be done by a limited set of enzymes [1, 4].

For the bottom-up construction of synthetic cells, there is a need for simple and modular ATP-generating pathways. In this context, the arginine deiminase (ADI) pathway is particularly attractive because of its simplicity and potential ease of implementation. It comprises the *arcA*, *arcB*, *arcC* and *arcD* genes, which encode three enzymes and an antiporter [1, 5, 6]. The arginine/ornithine antiporter ArcD renders the pathway effectively orthogonal to the rest of cellular metabolism by importing the substrate, arginine, while exporting the product, ornithine (Fig. 1). Within the cell, arginine deiminase ArcA converts arginine to citrulline plus NH_4_^+^. Ornithine carbamoyltransferase ArcB then converts citrulline plus inorganic phosphate into ornithine plus carbamoyl phosphate, and carbamate kinase ArcC converts carbamoyl phosphate plus ADP into ATP, CO_2_ plus NH_4_^+^ (Fig. 1). Thus, the pathway generates one ATP per arginine through only three enzymatic reactions and a single substrate/product antiporter. CO_2_ and NH_3_ (in rapid equilibrium with NH_4_^+^) can leave the cell via passive diffusion [7].

**Figure 1.**
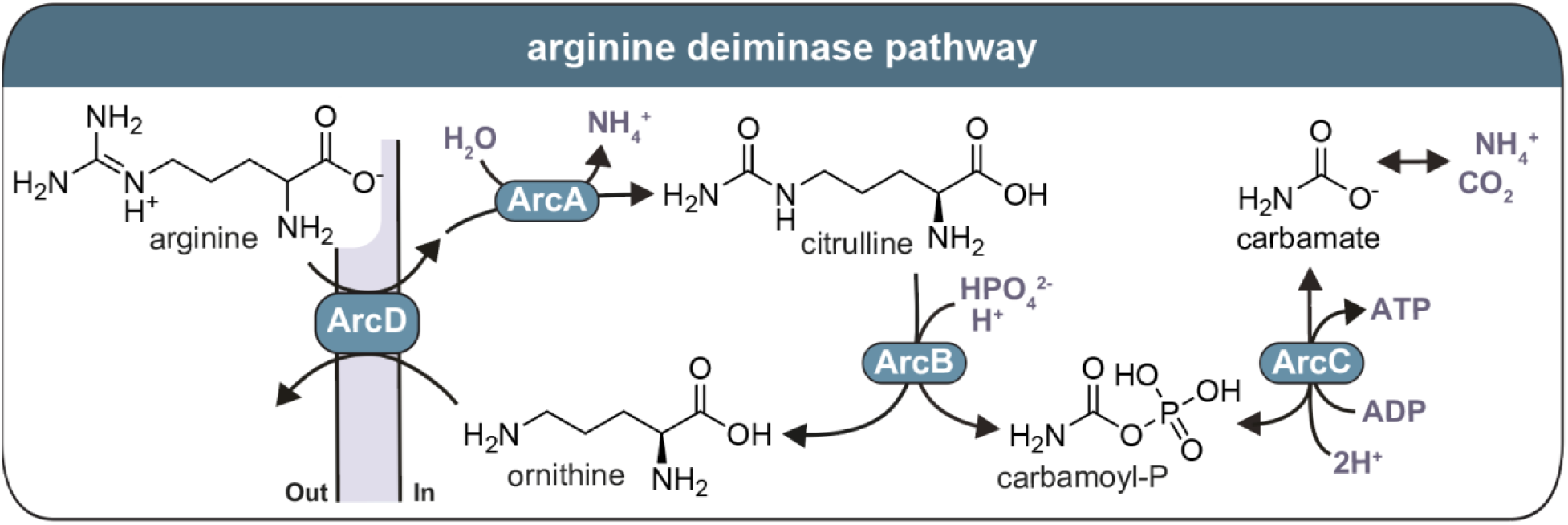
Schematic of the ADI pathway. ArcA: arginine deiminase, ArcB: ornithine carbamoyl-transferase, ArcC: carbamate kinase, ArcD: arginine/ornithine antiporter.

The ADI pathway is widespread in bacteria, archaea and some anaerobic eukaryotes [8], where it contributes both to energy conservation [9, 10] and to acid resistance. The overall pathway consumes net protons and thus alkalizes the cytoplasm [11–15]. Its potential as an engineered ATP-generating module has already been demonstrated in the slow-growing acetogen *Acetobacterium woodii*, in which expression of ADI genes from *Clostridium autoethanogenum* increased biomass yield, presumably through additional ATP production [16]. Likewise, implementing the pathway into mammalian cells has been shown to increase intracellular ATP levels [17]. However, in all naturally studied and engineered cases the pathway provided additional energy layered on top of native metabolism. Whether arginine breakdown can expand from supplementing cellular ATP production to being the sole pathway for ATP has not yet been shown.

Here, we address that question directly in *Escherichia coli* with the aim to select and evolve the pathway for full ATP production. Using a combinatorial ADI operon library, we first show that arginine deimination can function as an auxiliary ATP source. We then test whether the pathway can completely replace native ATP generation by constructing a strain in which ATP cannot be obtained from its main carbon source acetate, such that growth depends on the ADI pathway [18–21]. Although the pathway initially failed to restore growth, short-term laboratory evolution yielded a strain in which arginine breakdown became the sole source of ATP for cellular growth. Whole-genome sequencing revealed that this transition required inactivation of the principal electron-entry points into the respiratory chain. Finally, our results demonstrate that the ADI pathway, as initially designed and selected, is sufficiently active to sustain maintenance and growth in minimal medium of a living cell with a doubling time of ∼21 h and therefore likely to support the full ATP demands of a synthetic cell.

## 2. Materials and methods

### 2.1. Bacterial strains and culture conditions

The strains used in this study are listed in Suppl. Table 1. *E. coli* was routinely grown in LB medium (10 g/L tryptone, 10 g/L NaCl, 5 g/L yeast extract), with 15 g/L agar added for plates; 2x YP medium (16 g/L tryptone, 10 g/L yeast extract) was used for competent-cell preparation. Growth experiments used M9 minimal medium (3 g/L KH_2_PO_4_, 7.52 g/L Na_2_HPO_4_·2H_2_O, 0.5 g/L NaCl, 1 g/L NH_4_Cl), supplemented with 2 mM MgSO_4_ and 100 μM CaCl_2_, trace elements (134 μM EDTA, 31 μM FeCl_3_, 6.2 μM ZnCl_2_, 0.76 μM CuCl_2_, 0.42 μM CoCl_2_, 1.62 μM H_3_BO_3_, 0.081 μM MnCl_2_) and the relevant carbon source(s). Antibiotics were added as required (30 µg/mL chloramphenicol or 50 µg/mL kanamycin).

Plasmid construction (Suppl. Table 2) was done in *E. coli* DH10B. *E. coli* MG1655 was used to construct *E. coli* MG1655 Δ*atpBEFHAGD-atpC* and *E. coli* MG1655 *ΔcarB* strains. The KEIO collection was used as a template for the *ΔcarB* deletion cassette amplification [22].

### 2.2. Library design and assembly

Codon-harmonized gene variants were designed with the Codon Harmonizer tool [23] and ordered as gBlocks (IDT; Suppl. Table 3). Cloning PCRs used Q5 High-Fidelity 2x Master Mix (NEB) with 1 ng template, 500 nM of each primer and ddH_2_O to 50 µL (primers in Suppl. Table 4). Ribosome-binding site (RBS) variants were introduced as degenerate primer overhangs during amplification. To maximize the number of clones, the operon was assembled by Gibson assembly in two phases: two three-part assemblies first generated plasmids carrying *arcA*-*arcB* or *arcC*-*arcD*, after which the gene pairs were re-amplified and joined in a further three-part assembly to yield the complete *arcABCD* operon library. *E. coli* DH10B served as the cloning host for library construction; the assembled library was then isolated and used to transform *E. coli* BW25113.

### 2.3. Genomic deletions

Deletions were made by λ-Red recombineering [24]. To delete the *carB* gene in *E. coli* MG1655, the kanamycin antibiotic cassette from the Keio collection strain *E. coli* BW25113 *ΔcarB*, was amplified with primers flanking the gene (Suppl. Table 4). Using electroporation as described in Wenk et al. 2018 [20], this linear DNA fragment was introduced into *E. coli* MG1655 carrying the pSIJ8 plasmid, which expresses the λ­Red recombineering machinery (Suppl. Table 2). For this, *E. coli* MG1655 carrying the pSIJ8 plasmid was cultivated at 30 °C. Expression of the λ­Red recombineering machinery was induced by addition of 15 mM arabinose at OD_600_ ∼0.3 and incubation at 37 °C for 45 min. 1 mL of the cells were then harvested and washed 3 times with 10% (*v/v*) cold glycerol. Cells were electroporated with the linear DNA fragment (500 ng), recovered in SOB medium (20 g/L tryptone, 5 g/L yeast extract, 0.5 g/L NaCl, KCl 0.186 g/L) for 1 h at 37 °C and plated on LB plates containing kanamycin. Colony PCR was performed to confirm successful gene deletions. To delete the Δ*atpBEFHAGD-atpC* gene cluster in *E. coli* MG1655, the kanamycin antibiotic cassette was amplified from the pKD4 plasmid (Suppl. Table 2), with primers containing 50 bp homology arms that are flanking the *atp* cluster. Using this linear DNA fragment deletion was performed using λ­Red recombineering exactly as described above. Colony PCR was performed to confirm successful gene deletions.

### 2.4. Microplate growth assays

#### 2.4.1. Anaerobic growth

Prior to all growth experiments, the strains were streaked out from the glycerol stock on LB plates (supplemented with the respective antibiotic) and grown overnight at 37 °C. From these plates, precultures in M9 plus 20 mM glucose were inoculated for aerobic overnight growth at 37 °C. Precultures were washed three times in M9 and diluted to OD_600_ 0.1 in the relevant medium. In each well of a 96-well plate, 15 µL of diluted preculture was combined with 135 µL of medium and overlaid with 50 µL of mineral oil (Bio-Rad) to limit evaporation. Plates were incubated at 37 °C in a BioTek ELx800 absorbance reader (BioTek Instruments, USA) housed in an anaerobic tent, with a measurement cycle of 5 min fast linear shaking (19 Hz) followed by OD_600_ reading, for at least 24 h. Growth curves and doubling times of three biological and two technical replicates were extracted with an in-house MATLAB script [20].

#### 2.4.2. Aerobic growth

Precultures for aerobic growth experiments were prepared as in section 2.4.1. *Anaerobic growth*. The exact preculture medium composition can be found in Suppl. Table 5. The cells from precultures were harvested and washed three times with M9 and diluted to a final OD_600_ of 0.025 in the testing medium supplemented with carbon sources (Suppl. Table 5). For plate reader measurements, 150 μL of the culture was added to a Nunc 96-well microplate (Thermo Fisher Scientific) and covered with 50 μL of mineral oil (Sigma-Aldrich) to prevent evaporation. The growth experiments were performed in the BioTek Epoch2 plate reader (BioTek Instruments, USA). The mean and standard deviation of the three biological and two technical replicates is shown in the growth curves. For growth experiments in cell culture tubes, 5 mL of medium was inoculated with three times washed cells to an OD_600_ of 0.01 and the growth was measured using an Diluphotometer (Implen).

### 2.5. Cell growth and preparation for ornithine uptake experiments

Strains were streaked out from glycerol stocks on LB plates supplemented with chloramphenicol and incubated overnight at 37 °C. Precultures from single colonies grown in LB with chloramphenicol were 30x diluted (after growth for about 7 h) in M9 supplemented with 20 mM glucose, 40 mM arginine plus chloramphenicol and grown overnight at 37 °C. The next day fresh M9 media was inoculated with 1% preculture and grown to an OD_600_ of 0.5. Cells were harvested by centrifugation at 3000 x g for 10 min and washed two times with 20 mL of assay medium (100 mM KPi, pH 6.0 plus 10 mM glucose) to remove residual arginine. Cells were resuspended in assay medium to OD_600_ = 50 and assayed immediately or after overnight incubation on ice with or without loading with 1 mM ornithine.

### 2.6. Ornithine uptake assay

L-ornithine hydrochloride-[2,3-^3^H] (27Ci/mmol) was purchased from Moravek. The assay medium plus 1 µM ornithine (0.1 µM [^3^H]-ornithine + 0.9 µM unlabeled ornithine) (volume of 98 µL) was preheated for 2 min at 25 °C before 2 µL concentrated cell suspension was added (final OD_600_ = 0.5 to 1). At time intervals of 0, 10, 20, 30, 60 and 90 s the transport reaction was quenched with 2 mL ice-cold 0.1 M LiCl, and the mixture was filtered over prewetted nitrocellulose filters with a pore diameter of 0.45 µm (Protean, Cytiva). Subsequently, the filters were washed with 2 mL ice-cold 0.1 M LiCl and then dissolved in 2 mL Ultimagold^TM^ scintillation fluid (Perkin Elmer). Radioactivity, reflecting the uptake of [^3^H]-L-ornithine was determined with a Perkin Elmer Tri-carb 2800TR scintillation counter.

### 2.7. HPLC analyses

Amino acids and fermentation products were quantified by HPLC. End-of-fermentation samples (48 or 72 h) were derivatized with dabsyl chloride: 10 µL of clarified supernatant was mixed with 100 µL of 0.15 M NaHCO_3_ (pH 9.0), 200 µL of freshly prepared dabsyl chloride solution (1.3 g/L in acetonitrile) was added, and the mixture was incubated at 70 °C for 20 min, cooled, and diluted with 690 µL of 70% ethanol. Standards of arginine, citrulline and ornithine (2.5-20 mM) were treated identically, with norleucine as internal standard. Derivatized amino acids were separated on a Shimadzu LC 2030 C Plus with an Agilent Poroshell 120 EC-C18 column (250 × 4.6 mm, 4 µm; 690970-902) at 1 mL/min. A 30-min gradient ran from 25% eluent A (10 mM formic acid in water) and 75% eluent B (10 mM formic acid in acetonitrile), after which the column was re-equilibrated at 75/25 for 10 min; detection was at 436 nm.

### 2.8. Sequencing

ADI operons were sequenced by Sanger Sequencing from plasmids isolated with the GeneJET Plasmid MiniPrep Kit (Thermo Fisher) using four primers that anneal irrespective of gene variant (Suppl. Table 4; Macrogen Europe), and reads were aligned in Benchling. For evolved strains, whole-genome and plasmid sequencing were performed to identify adaptive mutations. For whole plasmid sequencing, plasmids were isolated using the NucleoSpin® Plasmid kit (Macherey-Nagel) and sequenced by Plasmidsaurus. For whole genome sequencing, genomic DNA was isolated using the Quick-DNA Fungal Bacterial Miniprep kit (Zymo Research) and sequenced by Illumina paired-end sequencing (150 bp reads) at Novogene. Plasmidsaurus-derived sequencing data was analysed using SnapGene and Novogene-derived data was analysed via breseq [25]. Whole genome sequencing data was deposited in the European Nucleotide Archive (ENA project ID: PRJEB129438).

### 2.9. Genome-scale metabolic modelling

The genome-scale metabolic model (GEM) of E. coli, iJO1366 [26] was retrieved in SBML format level 3 version 1 from BiGG Models [27]. Model simulations were performed using COBRApy 0.32.1 [28] and Python 3.14. Flux balance analysis (FBA) [29] was used as the mathematical framework to simulate growth on the assessed experimental conditions: growth on glucose; growth on glucose and arginine; growth on glucose with ADI; growth on glucose and arginine with ADI; growth on acetate and arginine with ADI, and growth on acetate and arginine with ADI in the evolved strain in anaerobic conditions. For the first four scenarios, glucose uptake rate (’EX_glc D_e’) was constrained to 10 mmol gDW^−1^ h^−1^ (with and without the ADI pathway), and in the conditions with glucose and arginine, arginine uptake rate (’EX_arg L_e’) was constrained to 20 mmol gDW^−1^ h^−1^ (with and without ADI pathway). To have the complete ADI pathway in the model, we added three reactions: Arginine deiminase (’argA’), Citrulline transport (’TCITRe’), and Citrulline exchange reaction (’EX_citr L_e’). To simulate growth on acetate and arginine, we set the glucose uptake to 0 and constrained the uptake of acetate (‘EX_AC_e’) and the uptake of arginine to a minimum of 5 mmol gDW^−1^ h^−1^ and a maximum uptake rate of 20 mmol gDW^−1^ h^−1^. In both scenarios, we added the ADI pathway and set to 0 the flux through the ATP synthase reaction (‘ATPS4rpp’). To simulate the evolved strain, we included the knockouts of the observed gene mutations: *Dld* (‘b2133’), *sdhA* (‘b0723’) and *nuoG* (‘b2283’). For all conditions, the oxygen exchange reaction (’EX_O2_e’) was set to 0 to simulate anaerobic conditions, and the lower bound of the ATP maintenance (’ATPM’) reaction was constrained to 8.4 mmol gDW^−1^ h^−1^. Under every scenario and described constraints, the biomass synthesis reaction (‘BIOMASS_Ec_iJO1366_WT_53p95M’) was defined as the objective function, and fluxes were computed using FBA. The ratio between the predicted maximum growth rate (h^−1^) and the glucose uptake rate (mmol gDW^−1^ h^−1^) was used to calculate the biomass yield in gDW mol^−1^ of glucose per condition. Additionally, Flux variability analysis (FVA) was run to compute the minimum and maximum production rate (mmol gDW ^−1^ h^−1^) of ornithine (’EX_orn_e’), citrulline (’EX_citr L_e’), glutamate (‘EX_glu_L_e’), acetate (’EX_ac_e’), and formate (‘EX_for_e’) at 90% optimality in the scenarios with glucose/glucose and arginine with and without the ADI pathway. The corresponding code and the model used to run the simulations are available in the following git repository: https://github.com/benit005/ADI_pathway.

## 3. Results

### 3.1. Construction of a combinatorial ADI operon library

To obtain a functional ADI pathway in *E. coli* without prior knowledge of the best combination of genes, we built a combinatorial operon library spanning a range of enzyme and coding variants from different organism and expression levels. Eleven gene variants were included: the wild-type *Lactococcus lactis arcA*, *arcB*, *arcC* and *arcD2* as well as their codon-harmonized versions for *E. coli*, the codon-harmonized *arcB* from *Pseudomonas aeruginosa* and the *arcD* from *Lactobacillus sakei* (higher turnover number than ArcD2 from *L. lactis* [30]) along with its codon-harmonized version. For each gene, a set of six ribosome-binding sites was designed with degenerate bases using the RedLibs algorithm [31, 32], which yields a small set of RBSs spanning a wide and approximately linear range of predicted translation-initiation rates. Combining 11 gene variants with six RBS variants per gene gives a theoretical library size of 62,208 (Fig. 2). The pADI library (*pADI_lib*) was transformed into *E. coli* BW25113. Proper assembly and library diversity were verified by colony PCR (Suppl. Fig. 1) and subsequent sequencing of 4 randomly selected colonies (Suppl. Table 6).

**Figure 2.**
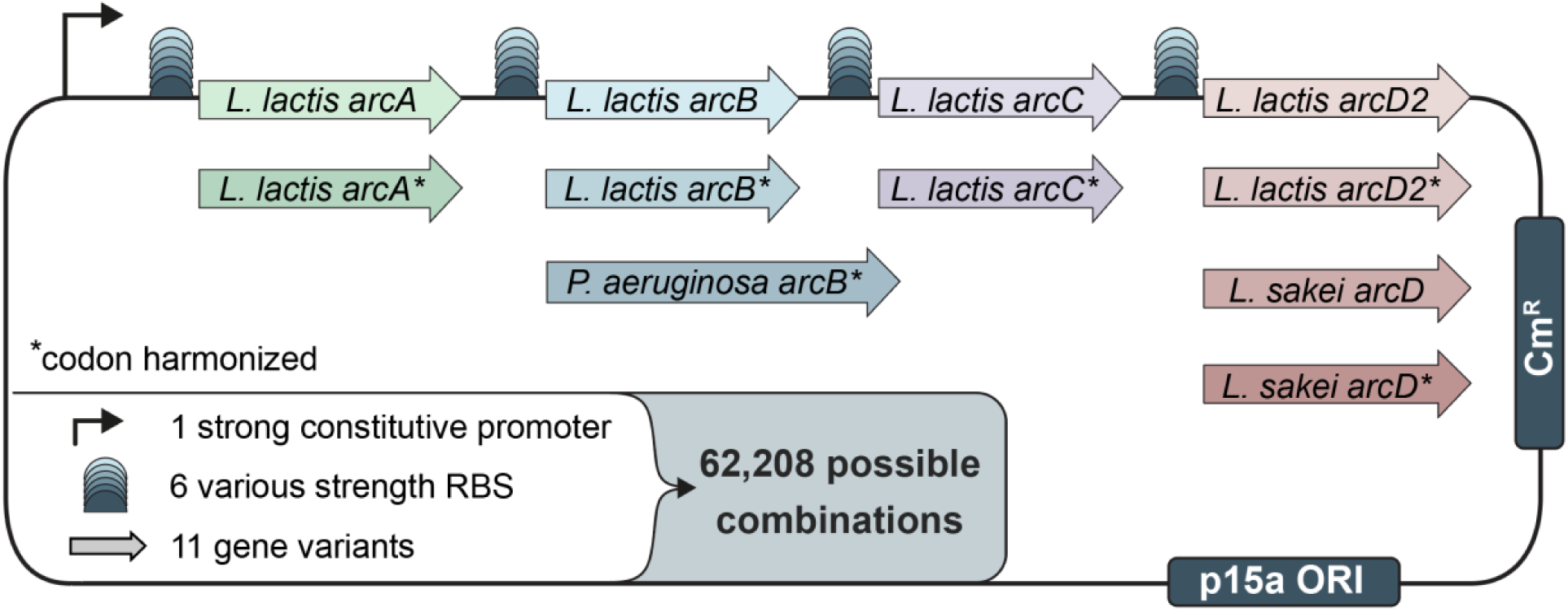
Design of the ADI-operon library. For each gene the source organism is shown, with an asterisk marking the codon-harmonized version; each gene is preceded by a RedLibs-designed degenerate RBS giving six variants of differing predicted translation-initiation strength. Operon expression was controlled by a strong constitutive promoter [20]. The plasmid contained the p15a origin of replication (ORI) and a chloramphenicol resistance gene (Cm^R^).

### 3.2. Functional ADI operons support auxiliary ATP generation

To identify active pathway variants, we screened the library under partial ATP limitation. Single *E. coli* BW25113 *pADI_lib* colonies were grown aerobically on glucose, then transferred to anaerobic growth in M9 with glucose plus arginine. During anaerobic fermentation in *E. coli*, ATP can only be obtained via substrate-level phosphorylation. This occurs mainly in glycolysis, as well as through acetate kinase when part of the carbon source is converted to acetate [33]. A cell with a functional ADI operon can obtain additional ATP by converting arginine to ornithine, CO_2_ and NH₄⁺. Of 94 colonies tested, 84 grew over 72 h with doubling times of 1.2 to 1.8 h, but maximum biomass varied widely from OD_600_ ∼0.1 to 2.0 (mean 0.62; Suppl. Fig. 2). Fifty-five strains secreted the pathway product ornithine (∼0.4 to 21 mM), and 49 also secreted the intermediate citrulline (0.1 to ∼37 mM). Because citrulline is the substrate of the ornithine carbamoyltransferase, its export indicates that at least the arginine deiminase and the antiporter are active. It also implies that the ornithine carbamoyltransferase or carbamate kinase is insufficiently expressed or active, so that little or no ATP is generated in these cells (Fig. 1). Strikingly, the five strains that produced the most ornithine (*pADI.9, pADI.11, pADI.39, pADI.54* and *pADI.83*) were also those that grew to the highest OD_600_, indicating that in these strains the pathway runs efficiently and the ATP it produces allows for extra biomass formation (Suppl. Fig. 2).

We next characterized the best clones in more detail. To this end, we sequenced plasmids of the top five and 17 randomly chosen strains (Suppl. Table 7). All gene variants, except the *L. lactis arcD2* and its codon-harmonized version were represented, and no single variant or RBS pattern clearly correlated with the best performers, indicating that several gene combinations support a functional pathway. The top five strains were further characterized in growth experiments on glucose either with or without arginine. Arginine significantly increased the final OD_600_ of all five ADI strains but had no effect on the empty-plasmid control (Suppl. Fig. 3). For the same strains ornithine and citrulline production (Fig. 3A) as well as fermentation products were analyzed (Fig. 3B). Supernatant analysis showed that ornithine accumulated to higher concentrations than citrulline in all strains, indicating efficient conversion of arginine to ornithine and functional arginine/ornithine antiport (Fig. 3A). Fermentation analysis showed increased production of formate, and decreased production of lactate in the presence of arginine (Fig. 3B). The accumulation of formate suggests increased flux from pyruvate into formate and acetyl-CoA via pyruvate formate lyase (PFL) and a shift away from lactate fermentation. Lactate fermentation natively supports redox-balanced glycolytic ATP production, which is less needed when the ADI pathway is active, allowing more pyruvate to be rerouted via PFL to acetyl-CoA for extra biomass formation. We confirmed that arginine/ornithine antiport was dependent on heterologously expressed ArcD, rather than endogenous basic amino acid transporters of *E. coli* by radiolabeled amino acid uptake experiments (Suppl. Note 1 and Suppl. Fig. 4). From the screen, *pADI.83* ranked highest by both growth and ornithine production and was therefore selected as construct for all subsequent work.

**Figure 3.**
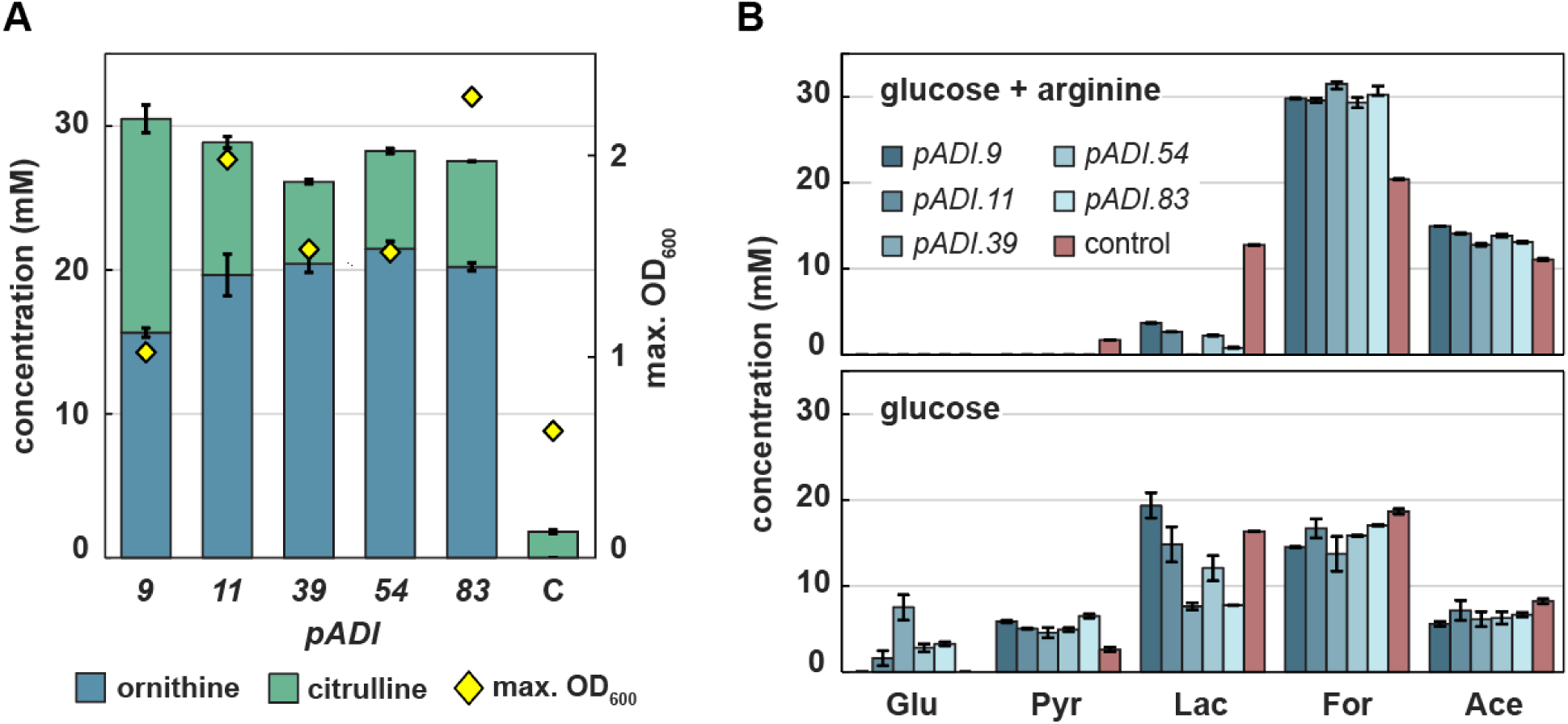
Characterization of the five best-performing ADI strains. (A) Citrulline and ornithine production by the strains grown in M9 medium supplemented with 20 mM glucose and 40 mM arginine (N=3). Error bars, s. d. of 3 biological replicates. (B) Fermentation products of the same strains on glucose with (upper panel) and without (lower panel) arginine (N=2). Bars indicate metabolite concentrations after 48 h of cultivation.

To test whether this yield increase is quantitatively consistent with ADI-derived ATP, we used the genome-scale metabolic model of *E. coli* (iJO1366) [26, 34] to simulate growth with and without the ADI pathway. Four conditions were compared: glucose alone and glucose plus arginine (in a 1:2 molar ratio matching the experiment), each with and without the pathway. The model predicted up to 1.4-fold higher biomass yield when arginine was present without the ADI pathway and up to a 3.2-fold higher biomass yield, when both the ADI pathway and arginine were present compared to glucose alone (Suppl. Fig. 5; git repository: https://github.com/benit005/ADI_pathway). In the other two cases the yield was unchanged. This closely matches the screen, in which the best strains reached 2.5-to 3.5-fold higher OD_600_ than the empty-plasmid control (Fig. 3A). As observed experimentally, the model also predicts a higher production of ornithine compared to citrulline for the best scenario (glucose and arginine with ADI). Together, the biomass increase, the fermentation shift and the model establish the ADI pathway as a functional auxiliary ATP-regenerating module in *E. coli*, extending to this organism the auxiliary role previously shown in engineered *Acetobacterium woodii* [16] and mammalian cells [17].

### 3.3. The upstream ADI pathway supplies carbamoyl phosphate *in vivo*

Having confirmed that the heterologous ADI pathway is fully functional in *E. coli*, we next tested stepwise whether it could replace *E. coli’*s native ATP metabolism. To this end, we constructed two strains whose growth is directly coupled to the breakdown of arginine via the ADI pathway. We established pathway activity in a modular manner: first the carbamoyl phosphate (CP)-generating module, using a CP-auxotroph (Fig. 4A), and then the complete pathway in an ATP-synthase deletion strain.

**Figure 4.**
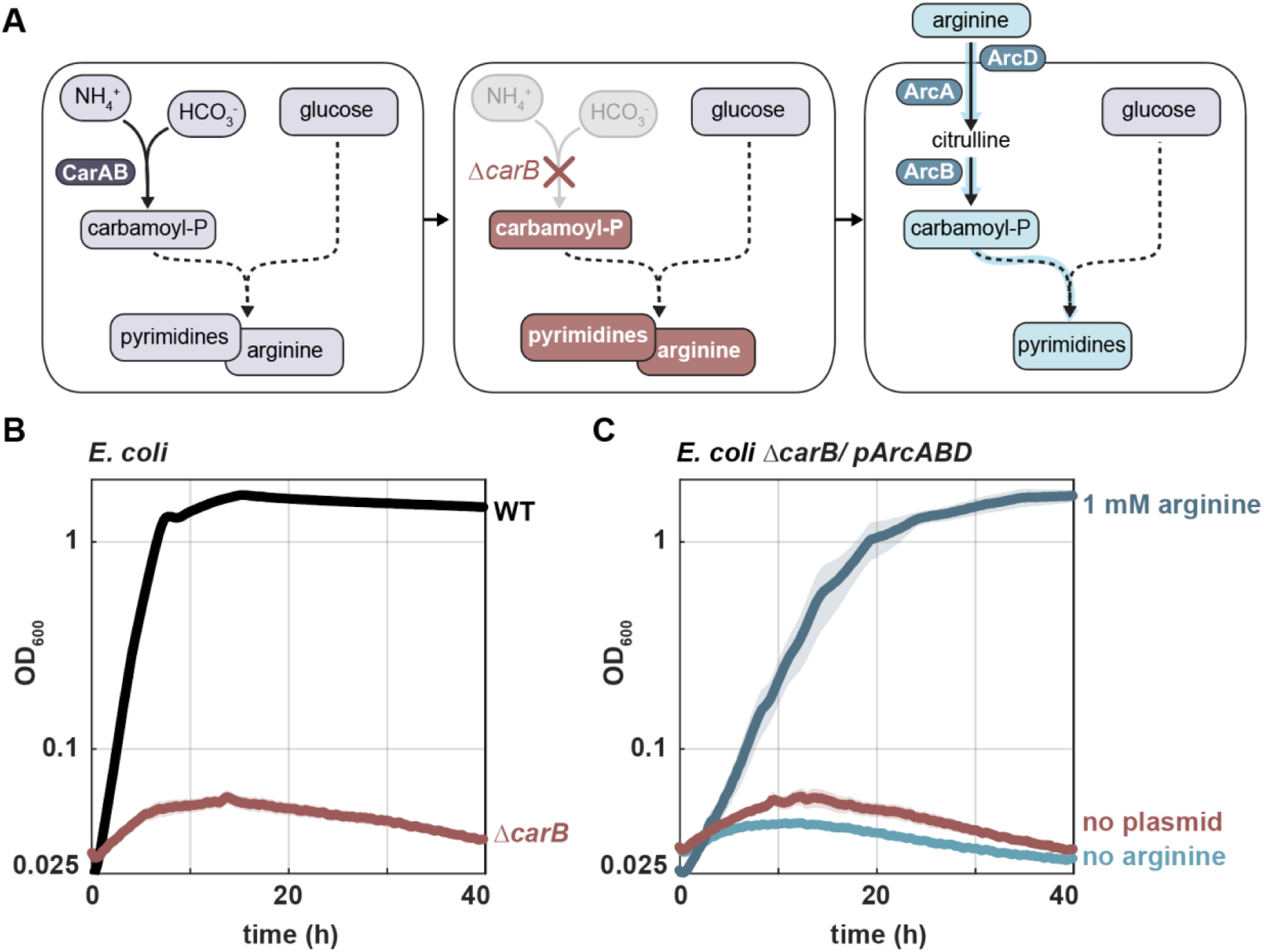
Testing the upper module of the ADI pathway in a CP-auxotroph selection strain. (A) CP-dependent biosynthesis of arginine and pyrimidine (left) is prevented when *carB* is deleted (middle). Expression of the *arcA*, *arcB* and *arcD* genes of the ADI pathway in the *E. coli* MG1655 *ΔcarB* strain and supplementation of arginine along with glucose should directly release the arginine auxotrophy and provide CP for pyrimidine biosynthesis. (B) Growth of the *E. coli* MG1655 wild type (WT, black) *versus* the *E. coli* MG1655 *ΔcarB* strain (red) on 10 mM glucose, (N=3). (C) *In vivo* activity of ArcA, ArcB and ArcD. The *ΔcarB/ pArcABD* strain grows on 10 mM glucose only when 1 mM arginine is supplied (dark blue). The no-plasmid and no-arginine controls do not grow (red and light blue, respectively), (N=3).

In *E. coli*, CP is essential for the biosynthesis of arginine and pyrimidines. It is produced by carbamoyl phosphate synthetase (CarAB), which forms carbamoyl phosphate from glutamine, bicarbonate and ATP. The small subunit (CarA) supplies glutamine-derived ammonia, and the large subunit (CarB) carries the synthetase activity. Deletion of the large subunit (Δ*carB*) renders *E. coli* auxotrophic for CP (Fig. 4B), and growth can be restored only by supplementing arginine together with uracil as a pyrimidine precursor (Suppl. Fig. 6). Expressing arginine deiminase and ornithine carbamoyltransferase along with the arginine/ornithine antiporter in this strain should relieve the CP auxotrophy in the presence of arginine. Indeed, transforming this strain with a *pADI.83*-derived plasmid carrying *arcA*, *arcB* and *arcD* but lacking *arcC* (*pArcABD*), allowed the strain to grow on glucose when arginine was supplied (Fig. 4C). In the absence of arginine, or of the plasmid, no growth was observed. ArcA, ArcB and ArcD are therefore sufficiently active *in vivo* to supply the CP required for growth, confirming that the upstream module of the pathway functions.

### 3.4. The ADI pathway makes auxiliary ATP but initially fails to serve as sole ATP source

Having confirmed that the upstream module is active, we next tested the full ADI pathway in an ATP depleted strain. To this end, we first made *E. coli’*s growth entirely dependent on substrate-level phosphorylation by deleting the entire ATP-synthase gene cluster (Δ*atpBEFHAGD-atpC*, hereafter Δ*atp*) (Fig. 5A). Next, we identified a carbon source on which ATP limitation would arrest its growth (Fig. 5B). Growth of the Δ*atp* strain on different carbon sources indicates to what extent its catabolism can yield ATP: the strain grew well on glucose (maximum OD₆₀₀ = 1.28, doubling time 1.4 h), to a lesser extent on glycerol (0.75, 2.8 h) and pyruvate (0.33, 2.6 h), and not at all on acetate. These results reflect that the first three carbon sources can still allow for a (decreasing) amount of ATP via substrate-level phosphorylation, whereas no substrate-level phosphorylation is possible from acetate. We therefore selected acetate as the carbon source for selection, since the Δ*atp* strain cannot make any ATP from it and therefore cannot grow on this substrate. Any growth must therefore be powered by an orthogonal ATP-producing pathway.

**Figure 5.**
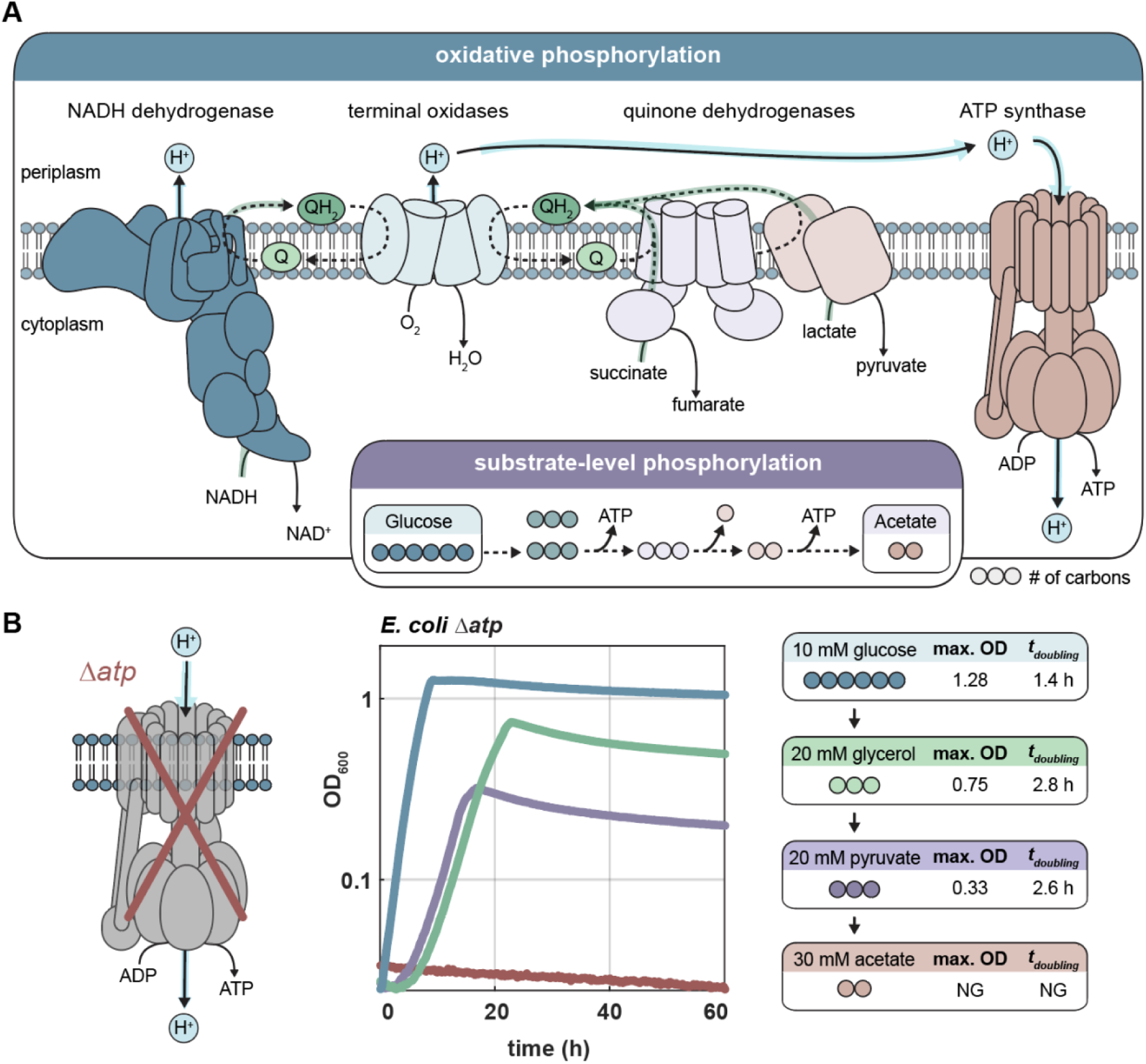
Construction of a chassis for growth-coupled selection of orthogonal ATP production. (A) Schematic of the respiratory chain and substrate-level phosphorylation. (B) Growth of the ATP-synthase mutant (*Δatp*) on different carbon sources, with maximum OD_600_ and doubling times indicated. Acetate does not support growth (NG) of the *Δatp* strain (N=3).

We then introduced *pADI.83* into the Δ*atp* strain and tested whether the ADI pathway could support growth on acetate and arginine. No growth was observed (Fig. 6A and Suppl. Fig. 7 showing a long-term measurement). Acetate is a particularly demanding substrate because it must first be activated to acetyl-CoA at the cost of one ATP (via acetate kinase and phosphate acetyltransferase) or two ATP (via the AMP-forming acetyl-CoA synthase) before it can enter the TCA cycle and biosynthesis [35]. In effect, activation of each acetate molecule consumes at least one ATP and therefore requires the conversion of at least one arginine.

**Figure 6.**
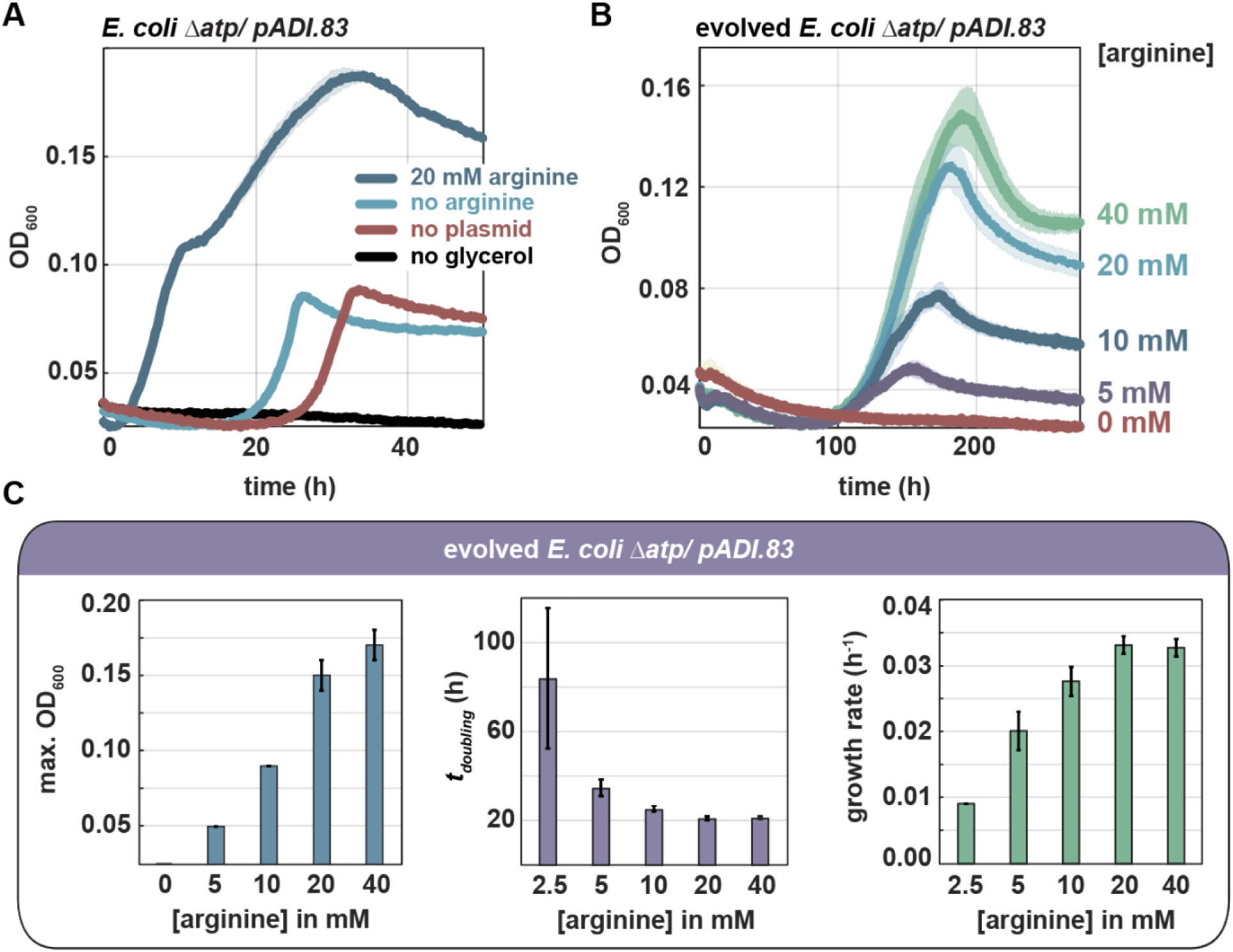
Growth phenotype of the *E. coli* MG1655 *Δatp* strain expressing the ADI pathway before and after evolution. (A) Glycerol-supported growth via the ADI pathway. *E. coli* MG1655 *Δatp/ pADI.83* was cultivated on 45 mM acetate with 1.5 mM glycerol and 20 mM arginine (dark blue). The no-plasmid (red) and no-arginine (light blue) controls don’t grow, indicating that the trace of glycerol supports but cannot by itself sustain growth. The no glycerol control shows no growth (black), (N=3). (B) Growth of the best performing evolved strain (evo1) on 45 mM acetate as a function of arginine concentration. Biomass (OD_600_) increases with the arginine concentration, and no growth occurs without arginine, (N=3). (C) Growth characteristic of the evolved strain, extracted from experiment in panel (B), and shown as a function of arginine concentration: (left) maximum OD_600_, (middle) doubling time, (right) growth rate.

To test whether the pathway was nonetheless active, we co-fed a small amount of glycerol as a limited source of ATP. On 45 mM acetate with 1.5 mM glycerol and 20 mM arginine, the strain grew to 2-fold higher OD_600_ than the no-plasmid and no-arginine controls with glycerol (Fig. 6A), confirming that arginine deimination is active and producing ATP *in vivo*, even though it is not yet sufficient to serve as the sole ATP source for growth.

### 3.5. Adaptive laboratory evolution enables complete replacement of native ATP generation

To remove the residual dependency on glycerol and force the cell to rely on arginine deimination alone, we performed short-term adaptive laboratory evolution on the *Δatp* strain plus *pADI.83*. Therefore, we serially passed the strain in M9 containing acetate, arginine and steadily decreasing concentrations of glycerol (from 2 mM to 0.5 mM) over approximately three weeks and 15 generations. We then isolated four evolved clones and tested them on acetate plus arginine in the absence of glycerol. The evolved strains grew on these two substrates alone reaching varying final OD_600_s (Suppl. Fig. 8). Further characterization of the best performing isolate (evo1) revealed that the biomass (reflected in OD_600_) increased with the arginine supplied (Fig. 6B). No growth occurred without arginine, while maximum OD_600_ increased with rising arginine concentrations to a maximum of ∼0.17 at 40 mM of arginine (Fig. 6C, left). The doubling times shortened to ∼21 h with 40 mM arginine (Fig. 6C, middle). Because acetate alone supports no growth in this background, and the biomass positively correlates with the arginine concentrations, growth must be driven by ATP derived from arginine deimination rather than by any residual host ATP production. These observations indicate that arginine deimination has become the sole ATP source supporting growth.

### 3.6. Loss-of-function of respiratory complexes enables ADI-dependent growth

To identify the genetic basis of growth via the ADI pathway as the sole source of ATP, we sequenced the genomes and plasmids of the parental strain and all four evolved strains. All mutations mapped to the genome, with none on the plasmid. Three of the identified mutations affected genes encoding respiratory dehydrogenases that feed electrons into the quinone pool: NuoG, an essential subunit of NADH:ubiquinone oxidoreductase (NDH-1, Complex I); Dld, the D-lactate dehydrogenase; and SdhA, the flavoprotein subunit of succinate dehydrogenase (Complex II), which also catalyzes the oxidation of succinate to fumarate in the TCA cycle (Suppl. Data 1). All three proteins were truncated: NuoG by a frameshift at His99 terminating at residue 117, SdhA at Tyr70, and Dld at Glu352. Notably, the Dld truncation was already present in the parental Δ*atp*/ *pADI.83* strain prior to any selection on acetate, while the SdhA truncation was found only in the best-performing evolved strain (evo1). This pattern suggests that Dld inactivation alone was insufficient to render arginine deimination the sole ATP source for growth, and that NuoG inactivation might have been the key adaptation. Subsequent loss of SdhA as the remaining electron-entry point might have created an additional fitness benefit. To confirm that these genomic mutations, rather than any undetected plasmid-level adaptation such as multimerization, were responsible for the phenotype, we cured the evolved strain of its plasmid and retransformed it with an unevolved plasmid. Both the retransformed derivative and the original evolved strain grew on acetate and arginine as sole carbon and energy sources (Suppl. Fig. 9, Fig. 6B), confirming that mutations supporting the ADI growth phenotype reside in the genome.

Metabolic model simulations on acetate and arginine with ATP synthase deleted and the ADI pathway active, sustained growth and showed zero flux through all the reactions affected by the corresponding gene mutations (see git repository: https://github.com/benit005/ADI_pathway). This suggests that the *nuoG*, *sdhA* and *dld* mutations are not detrimental to growth, as none of their reactions carries flux under optimal growth in the first place. Rather than having a metabolic flux-bearing role, their loss may instead reduce proton pumping into the periplasm, which, in the absence of ATP synthase, cannot be converted into ATP and may therefore be disadvantageous. Finally, consistent with the observation that no growth occurs without arginine, the model confirms that arginine deimination is the sole source of ATP supporting growth in this scenario.

**Figure 7.**
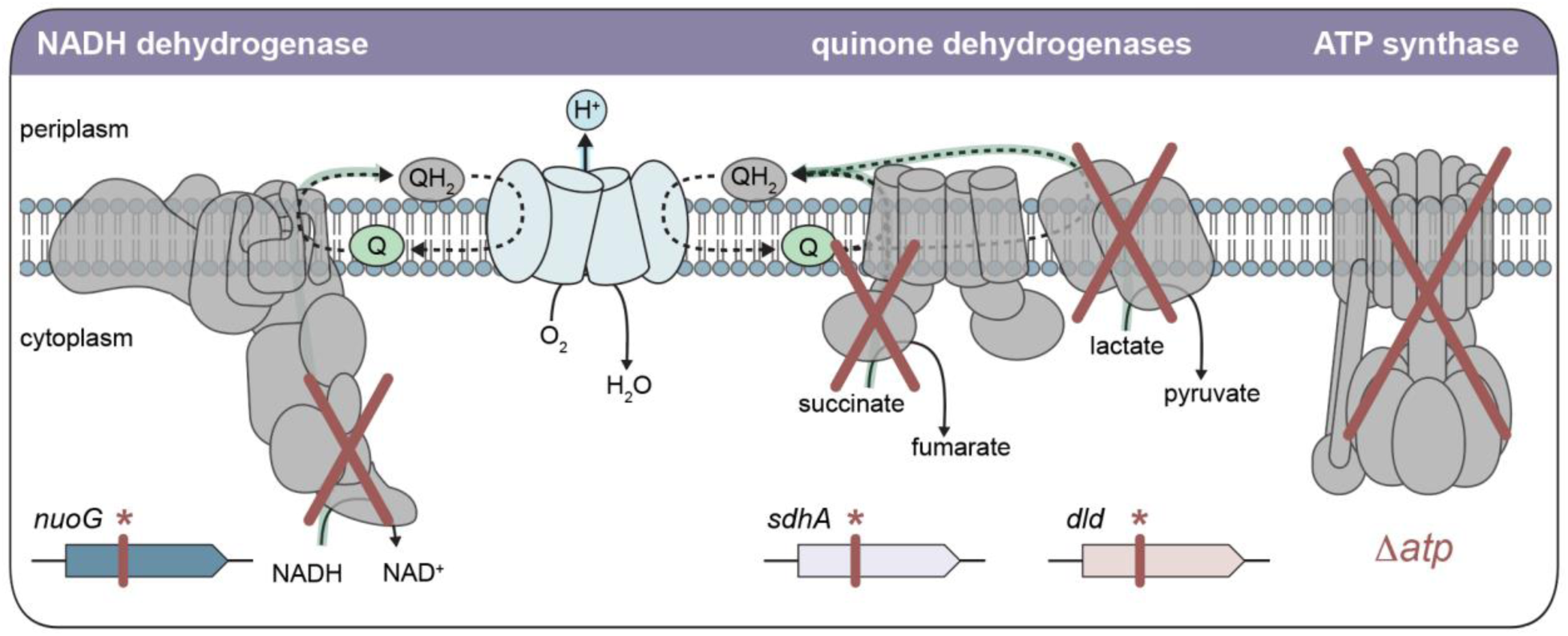
Mutations mapped onto the respiratory chain: truncations in *nuoG*, *sdhA* and *dld* inactivate the principal electron-entry points to the quinone pool in the *Δatp* background (see Table 1).

**Table 1.**
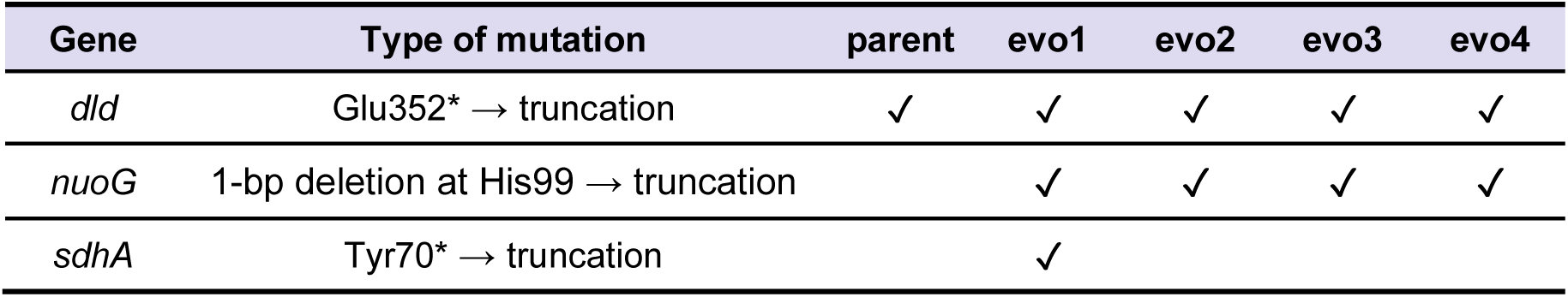
Respiratory chain mutations across evolved strains compared to parent strain. Whole genome sequencing data was deposited on ENA (Project ID: PRJEB129438).

| Gene | Type of mutation | parent | evo1 | evo2 | evo3 | evo4 |
|---|---|---|---|---|---|---|
| *dld* | Glu352* → truncation | ✓ | ✓ | ✓ | ✓ | ✓ |
| *nuoG* | 1-bp deletion at His99 → truncation |  | ✓ | ✓ | ✓ | ✓ |
| *sdhA* | Tyr70* → truncation |  | ✓ |  |  |  |

## 4. Discussion

We aimed to determine whether the native ATP metabolism of *E. coli* can be replaced by the minimal and orthogonal ADI pathway. When introduced as an auxiliary module, the pathway increased anaerobic biomass yield on glucose by approximately 3-fold, consistent with predictions from a genome-scale model and with previous demonstrations in *Acetobacterium woodii* [16]. However, under conditions in which native cellular ATP supply was eliminated, the ADI pathway alone was initially insufficient to support growth. Eventually, after short-term adaptive laboratory evolution in an *E. coli* ATP synthase mutant, arginine deimination became capable of sustaining growth as sole source of ATP. Importantly, the initially designed and selected ADI module was fully functional, and adaptation was required exclusively at the level of the host metabolism.

Several observations support the conclusion that growth of the evolved strains is driven by ATP derived from arginine deimination rather than from residual ATP production by the host. First, the ATP synthase mutant does not grow on acetate alone, providing a clean background in which acetate supplies carbon but no net ATP. Second, biomass formation scales with the concentration of arginine. Finally, curing the evolved strain of the plasmid results in a strain unable to grow on acetate and arginine, underlining the dependency on the plasmid (Suppl. Fig. 9). One could argue that acetate assimilation leads to ATP generation through succinyl-CoA synthetase in the TCA cycle. However, activation of acetate to acetyl-CoA consumes at least one ATP equivalent, rendering acetate metabolism effectively at best ATP neutral overall. Growth model simulations of the evolved strain further support these observations, confirming that ATP generated through the ADI pathway is required to sustain growth on acetate and arginine.

Adaptation to growth on acetate and arginine was supported by the inactivation of three respiratory dehydrogenases that normally feed electrons into the quinone pool. In cells lacking ATP synthase, electron transport can still drive proton pumping by NADH dehydrogenase and terminal oxidases, generating a proton-motive force that can no longer be discharged into ATP. The result is an energetically wasteful state in which reducing power is consumed and an electrochemical proton gradient is generated without sufficient metabolic return, possibly only serving a minor role in e.g. pH and ion homeostasis. Eliminating electron entry into the quinone pool prevents the formation of futile pumping of protons. The appearance of the *dld* mutation prior to selection on acetate and arginine suggests that loss of respiratory dehydrogenase activity is already advantageous in the ATP synthase mutant. Importantly, the parental strain carrying only the *dld* mutation remained unable to grow on acetate and arginine. Growth emerged only after additional mutation of *nuoG*, indicating that further disruption of respiratory chain activity is required before the ADI pathway can fully sustain cellular ATP demands.

Among the three mutations affecting the respiratory chain, the *sdhA* mutation is particularly informative because succinate dehydrogenase acts at the interface between respiration and central carbon metabolism. In addition to transferring electrons from succinate to the quinone pool, it catalyzes the oxidation of succinate to fumarate in the TCA cycle [36]. Loss of SdhA is therefore expected to eliminate flux through the oxidative branch of the TCA cycle. The TCA cycle normally serves to generate reducing power (e.g. NADH) for both respiration and biosynthesis. However, neither function is very useful here. Respiration is drastically reduced in the evolved strain, and the reduction state of acetate is similar to that of biomass, so additional NADH does not have a biosynthetic benefit. In addition, the TCA cycle can generate ATP via substrate-level phosphorylation through succinyl-CoA synthetase. However, this yields no net benefit for the same reason. Hence, TCA cycle flux would be not only wasteful but actively detrimental in this growth regime, as electron transfer via e.g. succinate dehydrogenase would over-reduce the quinone pool, generating a PMF that is not or only marginally used and may stall cellular processes.

The implications of these findings extend beyond microbial physiology to biotechnology and the construction of synthetic cells. Bottom-up synthetic cells require simple and robust mechanisms for ATP regeneration, and the ADI pathway is particularly attractive in this regard. It comprises only three enzymes and a single arginine/ornithine antiporter and has previously been reconstituted in proteoliposomes where it was demonstrated to generate ATP [30, 37]. Our demonstration that this pathway can serve as the sole ATP source for a growing and dividing cell extends this earlier work and strengthens the case for the ADI pathway as a promising energy-regeneration module for synthetic cells. The evolved strain doubled every ∼21 hours on acetate and arginine. Although substantially slower than wild-type *E. coli*, this result demonstrates that ATP generated exclusively through arginine deimination can support sustained cellular growth and division. Importantly, this was achieved under energetically demanding (minimal medium) conditions. In addition to the ATP demands for biosynthesis and cellular maintenance, growth on acetate as the sole carbon source places an additional energetic burden on the cell: acetate assimilation into central metabolism and the need to synthesize all organic components in the cell. Synthetic cells may instead be supplied with a richer medium with partially biosynthesized feedstocks, potentially reducing these energy demands and allowing for higher growth rates [1]. Growth on acetate and arginine can therefore be viewed as a stringent test of the capacity of the pathway to sustain cellular energy demands. Hence this work provides strong evidence that the ADI pathway is a suitable pathway for sustaining the full ATP demand of synthetic cells and likely allows their future growth at doubling times < 24 h. This answers the so far open question if the ADI pathway can supply ATP at a fast enough rate to support all cellular processes. The required evolutionary interruption of some respiratory chain complexes to get a fully functional ADI pathway in *E. coli* as seen in this work is not required for bottom-up synthetic cells as they will likely not be equipped with respiration complexes from the start.

Compared to glycolytic pathways, respiratory systems and photosystems, the ADI pathway offers practical advantages as an ATP-generation module for synthetic cells, as the alternatives are typically more complex to reconstitute bottom-up. Other widely used simple systems, such as the creatine-phosphate system do not support long-term ATP homeostasis, as ATP/ADP ratio already drops before the creatine-phosphate pool is depleted. Rhodopsin-based photosystems and artificial respiratory chains, though already reconstituted in liposomes, can only function together with an F_0_F_1_-ATP synthase [3], requiring functional expression of at least ten genes. Furthermore, rhodopsins and ATP synthase are both membrane-embedded, potentially occupying a large fraction of available membrane surface area, whereas the ADI pathway requires only a single antiporter. The growth demonstrated here confirms that sufficient membrane area remains available to support ArcD antiporter activity alongside the other membrane-embedded systems (e.g. transporters for other inorganic compounds) required for growth. Notably, early synthetic cells will likely strongly rely on membrane transporters as they will initially take up biosynthetic precursors from the medium rather than synthesizing them. We expect that for the ADI pathway, which only requires the ArcD antiporter, membrane occupancy in a synthetic cell will not be a major issue.

Overall, this work demonstrates for the first time that the ADI pathway is fully capable of sustaining cellular growth, underlining its potential as an energy module in synthetic cells. Future bottom-up synthetic cell efforts should therefore focus on expressing this pathway in liposomes. Next steps would include liposome-based transcription, translation and membrane insertion of the four ADI genes via cell-free protein expression systems such as PURE. Recent advances in co-translational membrane protein insertion in liposomes like the ProSecCO system, will be important for establishing functional expression of the ArcD antiporter [38]. Once established, the ADI pathway could support long-term ATP generation for PURE, eliminating the creatine-phosphate system currently limiting the functional lifetime of PURE-based transcription and translation. After establishment of the ADI encoded by PURE in liposomes, integration with other cellular functions such as lipid biosynthesis and DNA replication are obvious steps [39].

More broadly, an orthogonal ATP-generating module such as the ADI pathway could have applications in biotechnology, for example by increasing biomass yields or supporting the production of ATP-intensive compounds. A key limitation, however, is the relatively high cost of arginine as a substrate. Whether this cost is justified will depend on the value of the resulting product. For high-value products such as pharmaceutical compounds, increased productivity could potentially offset the cost of arginine. These considerations make the ADI pathway an interesting platform for exploring alternative strategies for ATP regeneration in both synthetic cell and biotechnological systems.

Lastly, the results obtained here also suggest that the ADI pathway in natural bacteria may in some environmental conditions also support full ATP production, if there is enough arginine. So far, its natural role is solely seen as an auxiliary role for ATP production or providing acid resistance. This work clearly provides the evidence that the kinetics of this pathway can at least fully support cellular ATP demand for maintenance and growth at high physiological capacity.

## Acknowledgements

The authors acknowledge the support of Sebastian Wenk for proof-reading the manuscript and providing plate readers and technical support. This publication is part of the project “Evolving life from non-life (EVOLF)” (SUMMIT.1.004) of the research program SUMMIT, which is financed by the Dutch Research Council (NWO) under the grant https://doi.org/10.61686/YLPDX93278. We acknowledge the support of the Dutch Research Council (NWO) via the Gravitation Project BaSyC (024.003.019) to J.v.d.O., B.P. and N.J.C., the Spinoza award (SPI 93-537) to J.v.d.O., and ERC Synergy (MetaDivide) to B.P. In addition, N.J.C. acknowledges support from his NWO Veni fellowship (VI.Veni.192.156) and ERC Starting Grant FASTFIX.

## Author contributions

K.S., J.H., N.J.C., B.P. and J.v.d.O. designed the study; K.S., J.H., Y.L., G.K.S., T.d.N., and Q.Y. performed the experimental work, E.v.A. and S.B.V performed the modelling experiments. K.S., J.H., N.J.C., J.v.d.O. and B.P. wrote and reviewed the manuscript.

## Data availability

The code and data required to reproduce the modelling results from the study are available at: https://github.com/benit005/ADI_pathway. The whole genome sequencing data was deposited on ENA (Project ID: PRJEB129438). Data and materials are available on request.

