## Supplementary Information for "Minimal, synthetic arginine deiminase pathway provides full ATP demand for cellular growth"

### Supplementary Figures

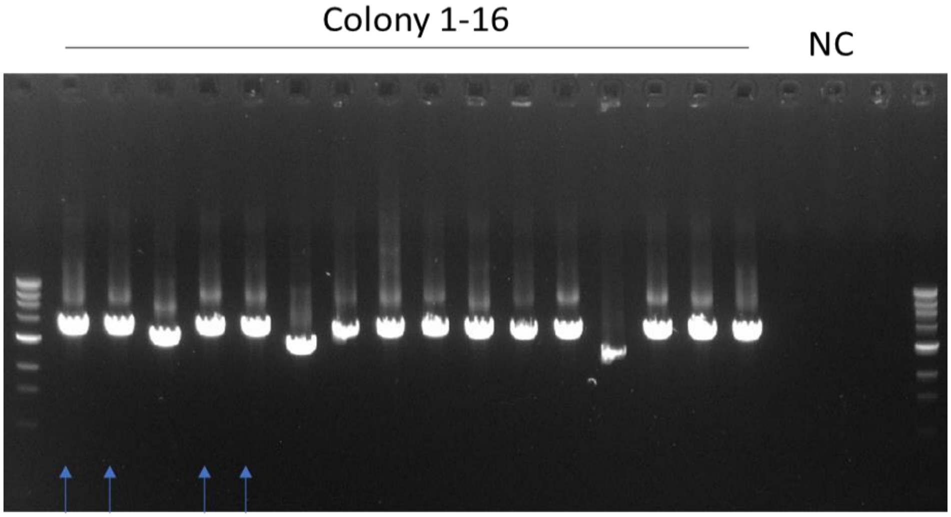


**Supplementary Figure 1.** Colony PCR on 16 colonies harboring the ADI-operon library plasmid. 13 of 16 colonies show the expected size. Colonies marked with blue arrows were used for sequencing of the operon variants.

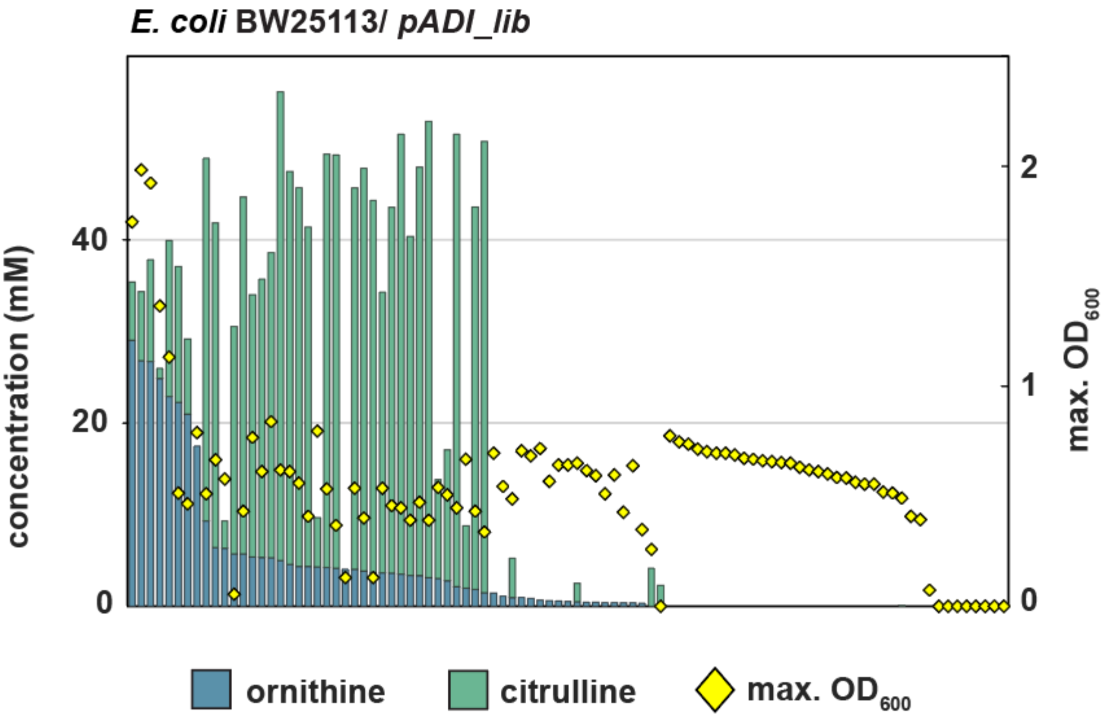


**Supplementary Figure 2.** Ornithine and citrulline production and maximum $OD_{600}$ of 94 ADI-library strains, sorted by ornithine production, then citrulline, and finally $OD_{600}$. The five highest ornithine producers also reach the highest $OD_{600}$.

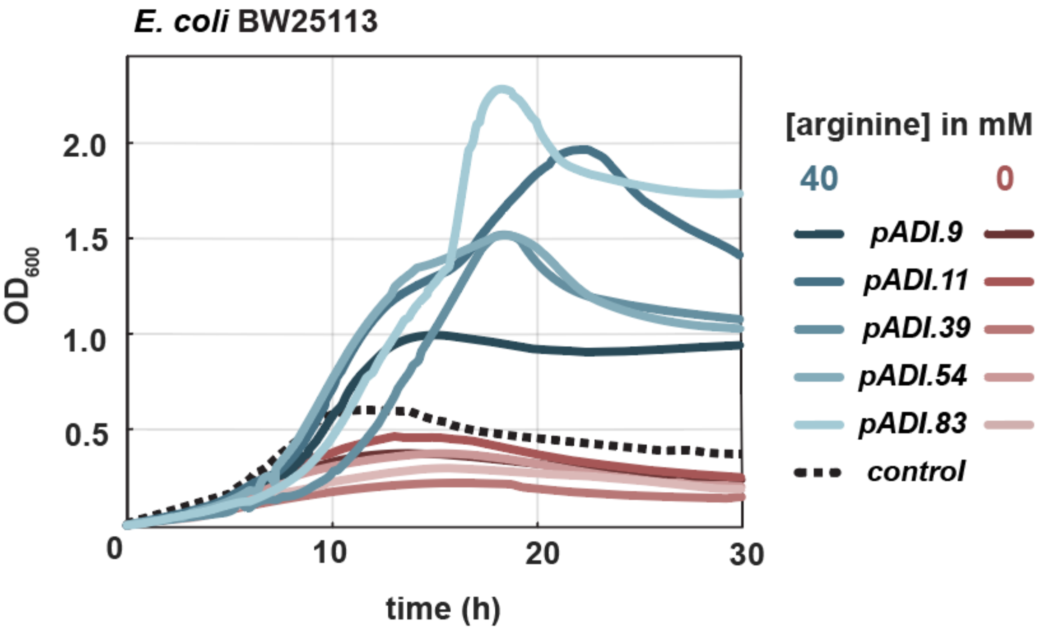


**Supplementary Figure 3.** Growth of the best-performing *E. coli* BW25113/ *pADI* strains on M9 supplemented with 20 mM glucose, with and without 40 mM arginine.

**Supplementary Note 1: Arginine/ornithine antiport via ArcD**

*E. coli* benefits from the ATP production by the synthetic arginine metabolic pathway (Fig. 3A), however arginine import and ornithine export may be mediated by *E. coli*'s endogenous transporters rather than the heterologous expressed ArcD from *L. sakei* (*Ls*ArcD). Supplementary Fig. 4A compares [$^3$H]-ornithine uptake (as a measure for *in vivo* activity of *Ls*ArcD) in strains with (*E. coli* BW25113/ *pADI.83*) and without ArcD (*E. coli* BW25113/ *pADI.83ΔarcD*). Both strains were either directly harvested or subjected to overnight starvation. [$^3$H]-ornithine uptake is significantly elevated in cells expressing *Ls*ArcD, indicating it being active and important for arginine/ornithine antiport. Overnight starvation of the cells reduced the transport activity by about 20%. The effect of varying intracellular ornithine concentration on [$^3$H]-ornithine uptake was assessed by overnight incubating the cells with ornithine. Suppl. Fig. 4B shows that overnight loading with ornithine does not increase the uptake rate of [$^3$H]-ornithine, most likely reflecting that the cells already have saturating levels of counter substrate to facilitate the exchange reaction. Suppl. Fig. 4C compares the uptake of [$^3$H]-ornithine in BW25113 harboring either of the plasmids: *pADI.9, pADI.11* or *pADI.83* or *pADI.83ΔarcD*. As anticipated from the strain design, the exchange activity is similar and, importantly, dependent of ArcD. Next, we determined the substrate specificity of ArcD by measuring [$^3$H]-ornithine/ornithine exchange in the presence of a 5000-fold excess of arginine, lysine and ornithine (Suppl. Fig. 4D). All these amino acids inhibit because they compete with [$^3$H]-ornithine for uptake. On the contrary, a 5000-fold excess of citrulline has little effect on the transport. Hence, we conclude that heterologous expressed ArcD facilitates not only ornithine/ornithine but also arginine/ornithine and lysine/ornithine exchange but very little or no citrulline/ornithine exchange, which is in line with studies on homologous proteins in *Lactobacillus brevis* ATCC 367 [1] and *L. lactis* [2]. Overall, our data show that the engineered *E. coli* cells take up arginine and excrete ornithine by the ArcD antiporter from *L. sakei* and they metabolize arginine and form ATP via ArcA and ArcC1 from *L. lactis* [3] and ArcB from *Pseudomonas aeruginosa*.

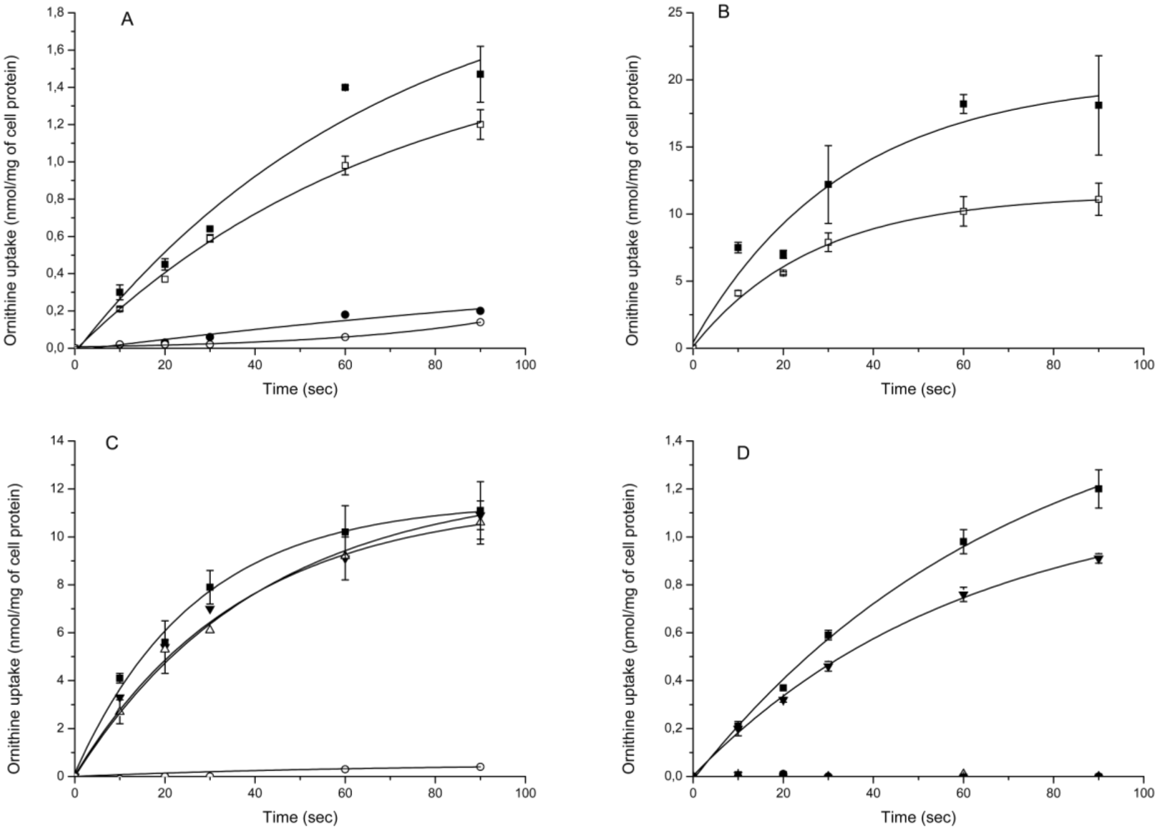


**Supplementary Figure 4.** Transport of [3H]-L-ornithine in engineered *E. coli* BW25113 strains. The basal assay medium was composed of 100 mM KPi, pH 6.0 plus 10 mM glucose, and the transport reactions were carried at a temperature of 25 °C. (A) Uptake of [3H]-L-ornithine (final concentration of 1 µM) in *E. coli* BW25113/ *pADI.83* (squares) and *pADI.83ΔarcD* (circles). The cells were used within 2 h after harvesting (closed symbols) or after overnight incubation (starvation) at 4 °C (open symbols). (B) Effect of overnight loading with 1 mM ornithine on the uptake [3H]-L-ornithine. Uptake of [3H]-L-ornithine was measured directly (final concentration 21 µM, closed squares) and after overnight incubation with 1 mM ornithine (final concentration 21 µM, open squares). (C) Uptake of [3H]-L-ornithine (final concentration of 21 µM ) in L-ornithine-loaded cells (see panel B) for *E. coli* BW25113 plus *pADI.9* (closed triangles), *pADI.11* (open triangles), *pADI.83* (closed squares), and *pADI.83ΔarcD* (open circles). (D) effect of 5 mM L-ornithine (filled circles), 5 mM L-arginine (open triangles), 5 mM L-citrulline (closed triangles), 5 mM L-lysine (closed stars) and no additions (filled squares) on the uptake of 1 µM [3H]-L-ornithine in overnight incubated cells of *E. coli* BW25113/ *pADI.83*; the cells were not preloaded with L-ornithine. All experiments: N=3.

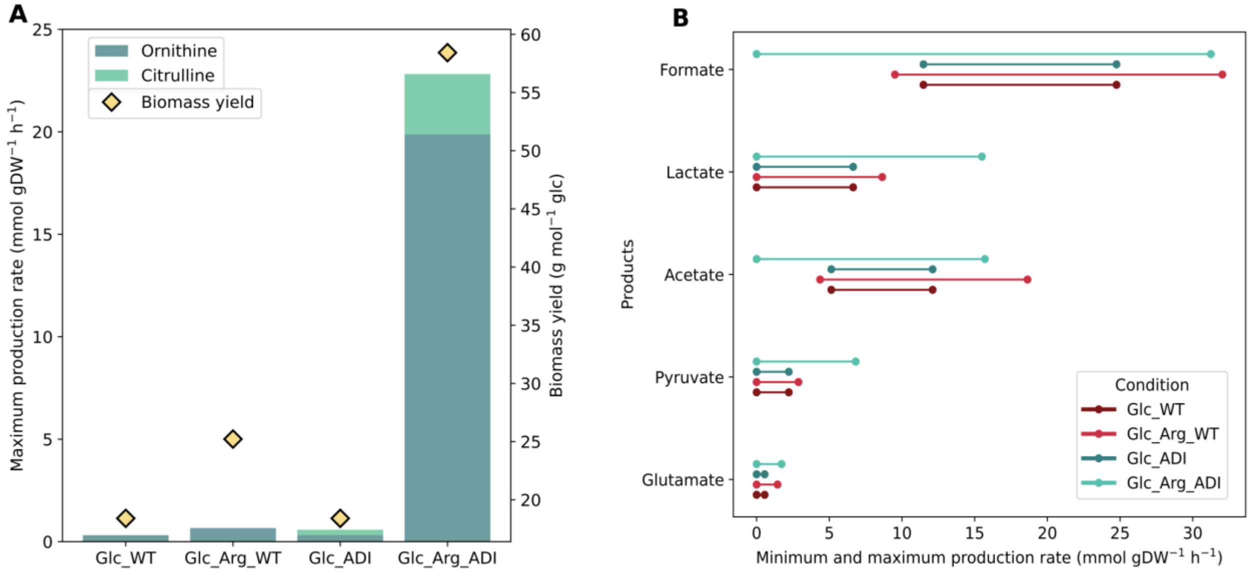


**Supplementary Figure 5.** (A) Biomass yields (g mol $^{-1}$ of glucose) and maximum production rates of ornithine and citrulline predicted by the *E. coli* model on glucose alone without the ADI pathway (Glc_WT, Glc_ADI, respectively) and on glucose plus arginine with and without the ADI pathway (Glc_Arg_WT, Glc_Arg_ADI, respectively). Biomass yields were obtained using FBA and the production rates correspond to the maximum rate predicted for ornithine and citrulline using FVA at 90% optimality. (B) Minimum and maximum production rate predicted for the specified products on glucose with/without the ADI pathway and on glucose plus arginine with and without the ADI pathway using FVA at 90% optimality.

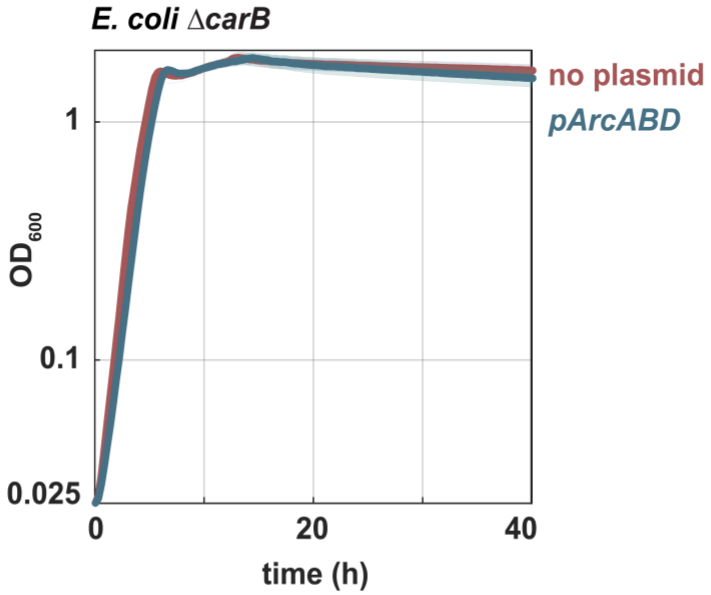


**Supplementary Figure 6.** Growth of the carbamoyl-phosphate auxotroph on 10 mM glucose supplemented with 1 mM arginine and 1 mM uracil (N=3).

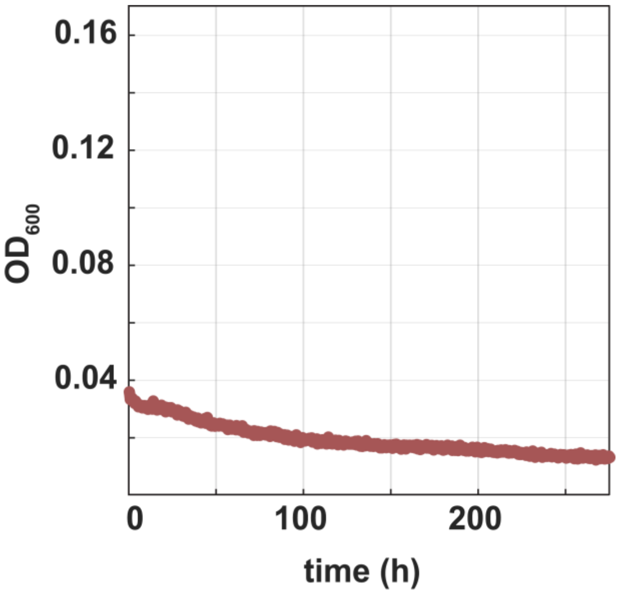


**Supplementary Figure 7.** Growth of *E. coli Δatpl pADI.83* on 45 mM acetate plus 20 mM arginine. No growth is observed over a long window of 250 hours (N=3).

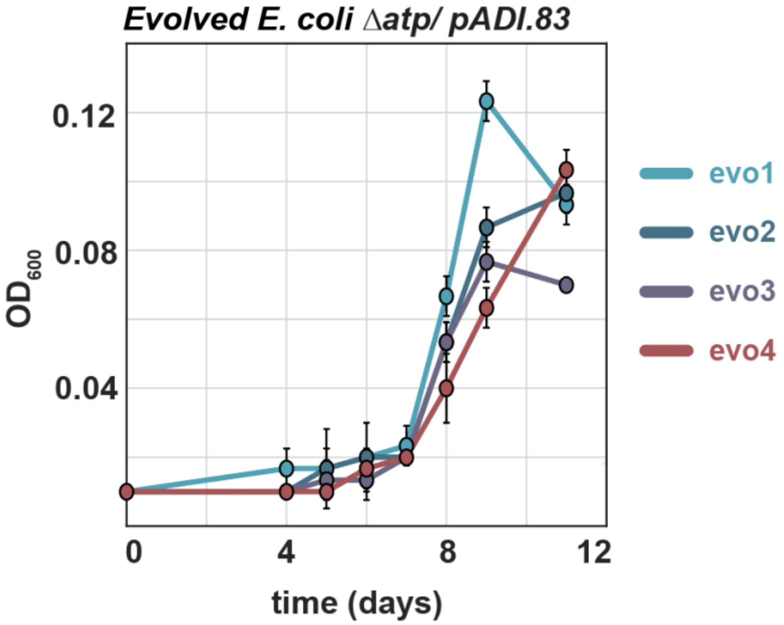


**Supplementary Figure 8.** Growth of four individual isolates of the evolved *E. coli Δatpl pADI.83* population in tubes on 45 mM acetate plus 20 mM arginine (N=3).

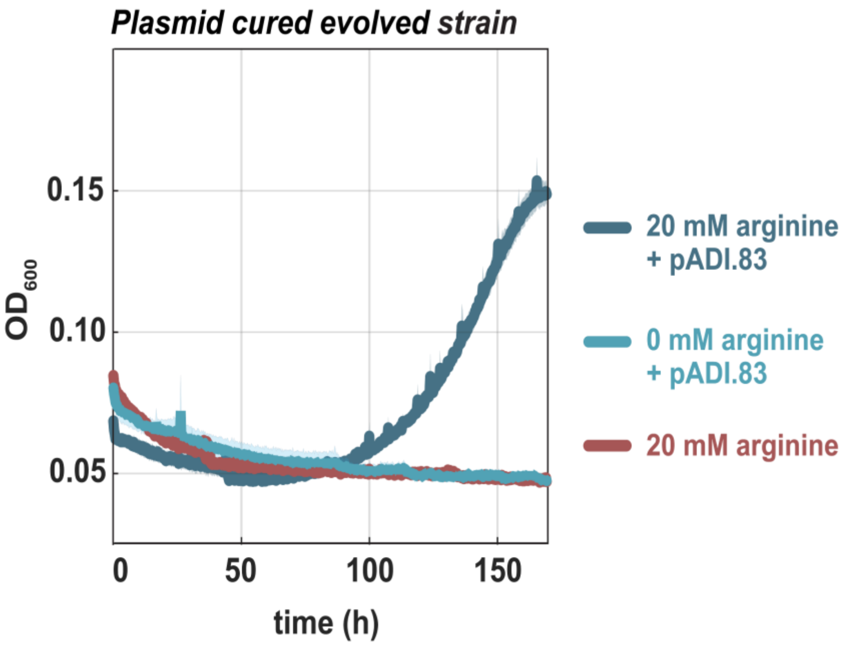


**Supplementary Figure 9.** Growth of the plasmid-cured evolved strain, retransformed with *pADI.83*, on 45 mM acetate with 20 mM arginine. Controls: 0 mM arginine (+ *pADI.83*) and 20 mM arginine (no plasmid) (N=3).

### Supplementary Tables

**Supplementary Table 1.** Strains used in this study.

| Strain | Use | Reference |
|---|---|---|
| *L. lactis* IL1403 | Source of ADI genes | [4] |
| *L. sakei* ATCC15521 | Cloning of the *LsArcD* | Gift from Jan Kok |
| *E. coli* BW25113 *ΔcarB* | Keio collection strain, used as *ΔcarB* template | [5] |
| *E. coli* DH10B | Cloning of the ADI library | [6] |
| *E. coli* BW25113 | Anaerobic assays with the ADI operon | [7] |
| *E. coli* MG1655 *ΔcarB (ΔcarB)* | Carbamoyl-phosphate auxotroph selection strain | This study |
| *E. coli* MG1655 *Δatp*BEFHAGD-*atp*C *(Δatp)* | ATP-synthase deletion; acetate-dependent selection strain | This study |

**Supplementary Table 2.** Plasmids used in this study.

| Plasmid | Use | Reference |
|---|---|---|
| *p*SIJ8 | Plasmid encoding the λ-Red recombineering machinery | [8] |
| *p*KD4 | Template for the deletion cassette containing the kanamycin-resistance gene | [7] |
| *pADI.9* | Plasmid encoding the ADI operon with encoding (*,[1]): *Ll*ArcA, h*Ll*ArcB, *Ll*ArcC1 and h*Ls*ArcD | This study |
| *pADI.11* | Plasmid encoding the ADI operon with encoding (*):: *Ll*ArcA, *Ll*ArcB, *Ll*ArcC1 and *Ls*ArcD | This study |
| *pADI.39* | Plasmid encoding the ADI operon with encoding (*):: h*Ll*ArcA, h*Pa*ArcB, *Ll*ArcC1 and *Ls*ArcD | This study |
| *pADI.54* | Plasmid encoding the ADI operon with encoding (*,[1]):: *Ll*ArcA, h*Ll*ArcB, *Ll*ArcC1 and h*Ls*ArcD | This study |
| *pADI.83* | Plasmid encoding the ADI operon with encoding (*):: *Ll*ArcA, h*Pa*ArcB, h*Ll*ArcC1 and h*Ls*ArcD | This study |
| *pArcABD* | *pADI.83*-derived plasmid lacking h*Ll*ArcC1 | This study |

(*) h = harmonized; *Ll*ArcA, *L. lactis arcA*; *Ll*ArcB, *L. lactis arcB*; *Pa*ArcB, *P. aeruginosa arcB*; *Ll*ArcC1, *L. lactis arcC1*; *Ls*ArcD, *L. sakei arcD*

([1]) *pADI.9* and *pADI.54* carry the same gene variants and differ only in RBS strength.

All *pADI* plasmids were based on a vector containing a p15a origin of replication and a chloramphenicol resistance gene. Transcription was initiated by a strong constitutive promoter [9], and translation was initiated by a set of RBSs spanning low to high translation-initiation rates (designed using the RedLibs algorithm [10, 11]).

**Supplementary Table 3.** G-blocks used in this study

| Name | Sequence |
|---|---|
| harmonized *L. Lactis arcA* (h*Ll*ArcA) | atgaataacggcattaacgtgaatagcgaaattggaaaactgaaaagcgtactcctccaccggccgggcgctgaagtggaaaacattacaccggacaccatgaaacagctgctgtttgatgatattccgtatctcaaaattgcgcagaaagagcatgatttctttgcgcagaccctgcgcgataacggcgccgaaaccgtgtatatagaaaacctggccaccgaagtgtttgaaaaaagctcggaaaccaaagaagaattcctgtcgcacctgctgcacgaagctggctaccgcccgggccggacctatgatggcctgaccgaatatctgaccagcatgagcaccaaagatatggtagaaaaaatatatgcgggcgtgcgcaaaaacgaactggatataaaacgcaccgccctgtcagatatggccggctcggatgccgaaaactatttctacctcaatccgctgccgaacgcttacttcacccgcgacccgcaggcgagcatgggcgtaggcatgacaattaacaaaatgacattcccggcgcgccagccggaatcactgattaccgagtatgtgatggcgaaccacccgcgcttcaaagataccccgatatggcgcgatcgcaatcatacaacccgcatagaaggcggcgatgaactgattctgaataaaaccaccgtagcgataggagtgagcgaacgcaccagctcgaaaaccattcagaacctggcgaaagagctgtttgccaatccgctgagcacctttgacaccgtactggctgtggaaattccgcataatcatgccatgatgcatctggataccgtatttaccatgattaatcatgatcagtttaccgtattcccgggcattatggatggcgcgggcaacataaacgtattcattctgcggccgggcaaagatgacgaagtggaaattgaacatctgaccgacctgaaagccgccctcaagaaagtactgaacctgagcgaactggacctgattgaatgcggcgccggcgatccgattgctgcgccgcgggaacagtggaacgatggcagcaacaccctggcgatagcccccggcgaaatagtgacctacgaccgcaattatgtaaccgtggagctgctgaaagagcatggcataaaagtgcatgaaattctgagctcagaactgggtcgtggacgtggtggtgctcgttgcatgagccagccgctgtggcgcgaagatctgtaa |
| harmonized *L. lactis arcB* (h*Ll*ArcB) | atgaccagcccgctgattaccaaagccgaagtaaatagcgtattccagggccggtcactgctggcggaaaaagatttcaccccggcggaaattaattacctggtggattttggcctgcatctgaaagccctgaaacagcagaatattccgcaccactacctggaaggcaaaaacattgctctgctctttgccaaaaccagcacccgcacccgcgcggcttttaccaccgcggccattgacctgggcgcgcatccggaatacctgggcgcgaatgatattcagctgggcattaaagaaagcaccgaagataccgcgcgggtactgggcagcatgtttgatgcgatagaacgccgcggcttttcgcagaaagaagtggaagatctggccaaatactcgggcgtgccggtgtggaacggcctgaccgatgattggcatccgacccagatgattgccgactttatgaccgtgaaagaaaattttggccatctgaaaggcctgaccctggtatacgtaggcgatggccgcaataatatggccaatagcctgatagtaaccggctcgatgctgggcgtaaacgtgcatattgtggcgccggatagcctgcatccgaccaaagaagtgatggatattgccaacaaatttgcggaaaaaagcggcgcgaaaccgctggccaccagcaacattgaagaaggcgtgaaagggccaatattatttacagcgacgtgtgggtaagcatgggcgaatcgaattgggaagaacgcgtgaaactgctgaccccgtaccgcataaccatggatatgctgaaaatgaccggcaacgcggacaacggccagctgatatttatgcactgtctgccggccttccatgataccgaaaccgaatatggcaaagaaataaaagaaaaatatggcctgaccgaaatggaagtaaccgacgaagtattccgctcgaaatatgcgcggcagtttgaagaagccgaaaaccgcatgcactcgataaaagccataatggcggccaccctgggcaacctgtttataccggccgtgccggaagattttaaataa |
| harmonized *L. lactis arcC1* (h*Ll*ArcC1) | atggtaaaacgcattgtggtagctctgggcggaaacgcgattctttcaaaagacccgagcgctaaggcgcagaaagaagccctggaagtgaccgcggaacagctgatgccgctgattaaagataacgaccgacagatggtggtaacccatggcaacggcccgcaagtgggcaatctgctgctgcagcagctcggcagcgattcggaaaaaaacccggcgatgccgctggataccgtgggcgccatgacacagggcgagattggcttttggacagtgcaggcctttacccgggccatggccaacaacgggctggagcattatccggtgattagcgccgtaacccagaccctggtggatgaaaacgatccggcgtttaaaaacgcgtcgaaaccgattggcccgttttacaccgaagaacagctgccggaaattaagaagcagtttccggattgggaactgatggaagacgcgggccggggctatcggcgggtagtgccgtcgccgaaaccgctggaaattattgaagctaaagcgtttggcccgatactggaatcgggctcgctcctgattgcgtcggggggcggcggcattcccgtaattaaagatggcaagggctatcagggcgtggaagctgtaattgataaagatttctcggccgcgaaacttgcggagatggtgggcgccgacgagctgctgattctgacctcaggcggcgacgtatatctgaattatggaaaagaaaatcaggtgcatctggaaaccgtaaaagtgtcggaaatgcggaaatatgtggacgaagaacggtttgccaaaggcagcatgggcccgaaagtagaagccgcggtatcgtttgtggaacggaccggcaagaccgcgaccattacctcactggacaacgtggaaaactttgtaaaatcgggctcgggcacccggattgtgccgtaa |

| | |
|---|---|
| harmonized *L. lactis arcD2*<br><br>(h*Ll*ArcD2) | atggaaaataagaaaaccaaaggaatttcactgtttgccctcctggccattattatttcgggcgccattggcggcggcgta<br>ttcaacctggcgaacgacctcgctaacggcagcacccccggcggcgtaatgattagctggctctttataggctttggaatt<br>ttcatgctggtactgtcgtttaaccggcttattaccattcggcccaaactgtcaggcgtatcggattatgcgcgggaaggatt<br>tggcgactttataggctttctcagcggctggggctactggattagcgcgtggaccggcaccattggctttgccgtgctgatg<br>atgacctcggcggactatttcttcccgtcaaaatttgcgaactcaaacggctcactgaccgtgctgagcgtaattattgtga<br>gcataataagctggatactgatgtttctcgtggaccggggcgtagaaaccgctgctctggtaaacgctgtggtgatgattg<br>ccaagctcattccgctggtggtatttagcataaccggaattatactctttaaagcggacgtattcacccagcatttttggcag<br>accttcacctcaaacctcgcggctgatggaagcgtaaaatcactcgtgtggcatgctatgaccgcgggcggcctcctgg<br>atcagattaaagggtcacttatggtaatggtatgggtatttgtaggaattgaaggggccgctatgatgggcaaccgagcc<br>aagaaaaaatcagataccgcgaaagctaccattattggcctcgccgtgctgctggtaatttatgtactgctgagcctgctg<br>ccgtatggctatatggaccaagcctcgctggccaacgtgaaagccccggggctggtgtatattctgaatgagatggtgg<br>gcgggtgggggggctcgcttatggccattggcctcatgatttcactgctgggcgcgtggctgtcatggaccatgctgccg<br>gtggaggctacccagcagctggccgagaaaaaactcctgccgtcgtggtttggcaagcttaacaaatatcatgctccgt<br>cgaactcgctgctgattacccaacttatgattcagatatttattataattacctattttgtagctaacgcgtataacgtatttattt<br>atatggccaccgcggtaattatgatttgctatgcgctggtgggcgcgtatctgctgaaaattggaattaaagaaggcagc<br>attaaaaacatactgataggctttttcacatttgcgtttcaggcgcttgctctgtatctgagcggctggcagtatgtgtggctg<br>gctatgattctgtatacaattggcttcattctgtttattaaagctaaaaaagaatcgcggcaggccataagcgtaaaagaat<br>gggccggcatgttcattgtagctctgctcggcgtgctggccattcttgtactgatttgcggggccaaaagcggcaccgcgc<br>tggatctgcgaggcctgctcgggtactaa |
| harmonized *L. sakei arcD*<br><br>(h*Ls*ArcD) | atgaccgaagaaaaaccggcgaaaaaaattggtctgttagcgctgattgcgttagtgattagcagctcaattggcagcg<br>gggtcttcggactgacctcagatctggccagcgcgtcagcgccaggtccggtactgatcgcgtgggtaattgtcggttttg<br>ggattctgatgctagcgctgtcattaaacaacctgttaatgaaagaaccagaactcgaagggattttctcatacgcggaa<br>aagggttttggtccctttgccggtttcattagcggttggggctattggttaagcgcgtggttagggaatgtagcgtttgccacc<br>attctgatgagcgccttaggctatttcttcccaatttttaaaagcgggcaaaatctgcccagcatcctggtggccagcgtgct<br>gtcatggtcactgacctattttgtgaaccgtggggtagaaggtgcggcggccatcaacaccttagtaaccatttgcaaatt<br>aattccgctctttgtcttcattatttttgggattgtactgtttaaaggccatctcttcacccaggcgttttggaacaacatgagca<br>gctcatttgtggccggtgatgtaatgtcacagatcaagaattgcatgatggtgatgatgtgggtctttgtcggcatcgaaggt<br>gcgagcatgctgtcagcccgtgccgaaaagaagtcagatgcgggcaaagcgaccattctgggactggtaagcctgct<br>cgccatttatatcctggcctcagtgctgccatatgggtatctgacccaggatcagctggcgtcaatcaagcagccggcca<br>tgctgtacatttttgaacaaatggtggggacgtgggggggttacttcatcggcgtggggctgatcatcagcattctggggg<br>cgtggctgtcatggaccatgctgccagccgaaaccatgttactgatggcgaagcagaacctgttaccggcgtacttcgg<br>ccgtgtcaataagaaaaaagcgccaacctttgcgctcgtcgtaaccgccgggttaattcaggtgttcctcttcacgctcct<br>gttcaccaccaaagcgtataatttcgcgtatagcctgtgcaccgcgagcattatcgtgtgctacatgttagtagcggcgtat<br>cagattaaatattcatggcccacctacaggaaaaaggaaaccgtcagcagctgctgattggggtgctggccctgctgt<br>ttgaaattgccggaatcctgatggccggcgtcagctacctgctgctttgctttatagcctatattccagggatttacttctatgg<br>ccgcgcccgcaaaaacaacgggcaccagcatttcctgagcaaaggggaatggctgatcaccaccattattgtcatcg<br>gcgcgattatcgggatttggctggtggtgagcgggaaaatcgtgatttag |
| harmonized *P. aeruginosa arcB*<br><br>(h*Pa*ArcB) | atggcttttaacatgcataaccgaaacctgttaagcctgatgcatcatagcacccgcgaactgcgatatctgttagatctgt<br>cacgcgatttaaaacgcgcgaaatataccggcaccgaacagcagcatctgaaacgcaaaaacatagcgctgattttt<br>gaaaagacctcaacacgcacccgctgcgcgtttgaagtggcggcgtatgatcagggcgcgaacgtgacctatattgat<br>ccgaactcatctcagattggccataaagaaagcatgaaagataccgcacgggtattaggccggatgtatgatgcgatt<br>gaatatcgcggctttaaacaggagattgtggaagagttagcaaaatttgcgggcgtaccggtgtttaacggcctgaccg<br>atgagtatcatccgacccagatgttagcggatgtgctgaccatgcgcgaacatagcgataaaccgctgcatgatattag<br>ctatgcgtatctgggcgatgcacgcaacaacatgggcaactcactgctgctgattggcgcgaaattaggcatggatgta<br>cgcattgcggctccgaaagcattatggccgcatgatgaatttgtggcgcagtgcaaaaaatttgcggaagagagcggc<br>gcgaaactgaccttaaccgaagacccgaaagaggcggtgaaaggcgtggattttgtgcataccgatgtgtgggtgtct<br>atgggcgaaccggtggaggcgtggggcgaacggattaaagagctgctgccatatcaggtgaacatggaaattatgaa<br>agcgaccggcaacccgcgggcgaaatttatgcattgcctgccggcatttcataacagcgaaaccaaagtgggcaaac |

| | |
|---|---|
| | agattgcggaacagtacccgaacctggcgaacggcattgaagtgaccgaagatgtgtttgaatctccatataacattgc gtttgaacaggcggaaaaccgcatgcacaccattaaagcaattttagtgtctaccttagcggatatttaa |

**Supplementary Table 4.** Primers used in this study.

| Primer Name | Primer Sequence (5’ to 3’) | Description |
|---|---|---|
| ArcA_fw | atgaacaatggaattaatgttaac | Amplification of *arcA* from *L. lactis* |
| ArcA_rv | ttacaaatcttcacgccaaag | |
| ArcB_fw | atgacatcaccacttattacaaaag | Amplification of *arcB* from *L. lactis* |
| ArcB_rv | ttatttaaaatcttcaggaactgc | |
| ArcC1_fw | atggtaaaacgtattgttgtagc | Amplification of *arcC* from *L. lactis* |
| ArcC1_rv | ttaaggaacaattcttgttcc | |
| ArcD2_fw | atggaaaacaagaaaacaaaagg | Amplification of *arcD* from *L. lactis* |
| ArcD2_rv | ttagtagcctagtaatccccg | |
| LL_arcA_H_fw | atgaataacggcattaacgtg | Amplification of *L. lactis arcA* harmonized from g-block |
| LL_arcA_H_rv | agtccatgggttaggctatcttac | |
| LL_arcB_H_fw | atgaccagcccgctgattac | Amplification of *L. lactis arcB* harmonized from g-block |
| LL_arcB_H_rv | tgagtattggactcacaaacttatttaaaatc | |
| LL_arcC_H_fw | atggtaaaacgcattgtggtag | Amplification of *L. lactis arcC* harmonized from g-block |
| LL_arcC_H_rv | agttccggtgtcaccggtag | |
| LL_arcD_H_fw | atggaaaataagaaaaccaaagg | Amplification of *L. lactis arcD* harmonized from g-block |
| LL_arcD_H_rv | tcggcattagtacccgagc | |
| PA_ArcB_H_fw | atggcttttaacatgcataac | Amplification of *P. aeruginosa arcB* harmonized from g-block |
| PA_ArcB_H_rv | tgagtattggactcacaaacttaaatatc | |
| LS_arcD_H_fw | atgaccgaagaaaaaccgg | Amplification of *L. sakei arcD* harmonized from g-block |
| LS_arcD_H_rv | aagcagcaactgagtcggc | |
| LS_ArcD_fw | atgacggaagaaaaaccagc | Amplification of *arcD* from *L. sakei* |
| LS_ArcD_rv | ctaaataacgattttccctgaaacaac | |
| pACYC_ADI_fw | tgccgactcagttgctgc | Amplification of pACYC184 backbone |
| pACYC_ADI_rv | caaagttggacgattacgtc | |
| L. lactis ArcA RL_fw | gacgtaatcgtccaactttgdaarggagattaattaatgaa caatggaattaatgttaac | *L. lactis arcA* fw primer with RedLibs RBS |
| L. lactis ArcB RL_fw | atagcctaacccatggacttaacbggargtttaatatgaca tcaccacttattacaaaag | *L. lactis arcB* fw primer with RedLibs RBS |

| | | |
|---|---|---|
| L. lactis ArcC RL_fw | gtttgtgagtccaatactcatatgggbggktatttaatggtaaaacgtattgttgtagc | *L. lactis arcC* fw primer with RedLibs RBS |
| L. lactis ArcD RL_fw | ctaccggtgacaccggaacttygaggagtatttcdtatggaaaacaagaaaacaaaagg | *L. lactis arcD* fw primer with RedLibs RBS |
| L. lactis ArcA Harmonized RL_fw | gacgtaatcgtccaactttgaattaagggbgcaatwatgaataacggcattaacgtg | *L. lactis arcA* harmonized fw primer with RedLibs RBS |
| L. lactis ArcB Harmonized RL_fw | gatagcctaacccatggactctsaaggaggttghatatgaccagcccgctgattac | *L. lactis arcB* harmonized fw primer with RedLibs RBS |
| L. lactis ArcC Harmonized RL_fw | gtttgtgagtccaatactcactgktggaggtttgtdatggtaaaacgcattgtggtag | *L. lactis arcC* harmonized fw primer with RedLibs RBS |
| L. lactis ArcD Harmonized RL_fw | ctaccggtgacaccggaacttaahggmgcaattaatatggaaaataagaaaaccaaag | *L. lactis arcD* harmonized fw primer with RedLibs RBS |
| L. sakei ArcD RL_fw | ctaccggtgacaccggaacttaaggdagagyttaatatgacggaagaaaaaccagc | *L. sakei arcD* fw primer with RedLibs RBS |
| L. sakei ArcD Harmonized RL_fw | ctaccggtgacaccggaactagdtaaggaatawcctatgaccgaagaaaaaccgg | *L. sakei arcD* harmonized fw primer with RedLibs RBS |
| P. aeruginosa ArcB Harmonized RL_fw | gatagcctaacccatggactaattggvgmagatattatggcttttaacatgcataac | *P. aeruginosa arcB* harmonized fw primer with RedLibs RBS |
| pTN03_fw | tataggatcctaactcgagc | Backbone primers used in plasmid for 2-step assembly |
| pTN03_rv | gccttaatacgttattagccg | |
| pTN03_fw_arcAB | gtttgtgagtccaatactcatataggatcctaactcgagc | Primers used in 2-step assembly of *arcA* and *arcB* |
| pTN03_rv_arcAB | caaagttggacgattacgtcgccttaatacgttattagccg | |
| pTN03_fw_arcCD | tgccgactcagttgctgctttataggatcctaactcgagc | Primers used in 2-step assembly of *arcC* and *arcD* |
| pTn03_rv_arcCD | tgagtattggactcacaaacgccttaatacgttattagccg | |
| carB_KO_fw | ctgcaaacctgcgtgtcac | Used to amplify *carB*::Km KO cassette from Keio collection strain. Also used to verify *carB* deletion. |
| carB_KO_rv | gtatgcaacttcaccgtcgataagtg | |
| atp_KO_fw | gttttatacgacacgcggcatacctcgaagggagcaggagtgaaaaacgtgatggtgtaggctggagctgc | Used to amplify Δ*atpBEFHAGD-atpC*::Km KO cassette from *pKD4* plasmid. |
| atp_KO_rv | gccagcctgtttccagactggcttttgtgcttttcaagccggtgttacatgtccatatgaatatcctccttagttcc | |
| atp_V_fw | gatctgaagcacgctttatcac | Use to verify *atp* deletion. |
| atp_V_rv | atcagcaggatctatgtgaacg | |

**Supplementary Table 5.** Carbon sources used in aerobic growth experiments.

| Strain | Relaxing Medium | Testing Medium |
|---|---|---|
| *E. coli* MG1655 *ΔcarB/ pADI.83* | 10 mM glucose<br>1 mM arginine<br>1 mM uracil | 10 mM glucose<br>1 mM arginine |
| *E. coli* MG1655 Δ*atp/ pADI.83* | 10 mM glucose | 45 mM acetate<br>1.5 mM glycerol<br>20 mM arginine<br>2.5 mM $MgCl_2$ |
| Evolved *E. coli* MG1655 Δ*atp/ pArcABD* | 10 mM glucose | 45 mM acetate<br>20 mM arginine<br>2.5 mM $MgCl_2$ |

**Supplementary Table 6.** Gene variants in the operon of the four randomly sequenced colonies after library assembly. h = harmonized; *Ll*ArcA, *L. lactis arcA*; *Ll*ArcB, *L. lactis arcB*; *Pa*ArcB, *P. aeruginosa arcB*; *Ll*ArcC1, *L. lactis arcC1*; *Ls*ArcD, *L. sakei arcD; Ll*ArcD2, *L. lactis arcD2.*

| **Colony** | ***arcA* variant** | ***arcB* variant** | ***arcC* variant** | ***arcD* variant** |
|---|---|---|---|---|
| **1** | h*Ll*ArcA | *Ll*ArcB | *Ll*ArcC1 | h*Ls*ArcD |
| **2** | *Ll*ArcA | h*Ll*ArcB | h*Ll*ArcC1 | *Ls*ArcD |
| **4** | h*Ll*ArcA | h*Pa*ArcB | *Ll*ArcC1 | *Ls*ArcD |
| **5** | h*Ll*ArcA | h*Ll*ArcB | *Ll*ArcC1 | *Ll*ArcD2 |

**Supplementary Table 7. Genes in ADI plasmids of sequenced colonies including predicted translation initiation rate (TIR) of RBS**. Missing or incomplete sequencing results are marked with a dash (-). 14 out of 22 sequenced plasmids were sequenced completely. The five strains that reached the highest biomass yield are indicated in red. h = harmonized; *Ll*ArcA, *L. lactis arcA*; *Ll*ArcB, *L. lactis arcB*; *Pa*ArcB, *P. aeruginosa arcB*; *Ll*ArcC1, *L. lactis arcC1*; *Ls*ArcD, *L. sakei arcD.*

| Colony | *arcA* TIR | *arcA* variant | *arcB* TIR | *arcB* variant | *arcC* TIR | *arcC* variant | *arcD* TIR | *arcD* variant |
|---|---|---|---|---|---|---|---|---|
| **1** | 0.909 | h*Ll*ArcA | 0.753 | h*Ll*ArcB | - | - | 0.239 | h*Ls*ArcD |
| **4** | 0.066 | *Ll*ArcA | 0.075 | h*Ll*ArcB | 0.918 | *Ll*ArcC1 | - | - |
| **9** | **0.229** | ***Ll*ArcA** | **0.600** | **h*Ll*ArcB** | **0.424** | ***Ll*ArcC1** | **0.756** | **h*Ls*ArcD** |
| **10** | 0.429 | *Ll*ArcA | 0.934 | *Ll*ArcB | 0.424 | *Ll*ArcC1 | 0.239 | h*Ls*ArcD |
| **11** | **0.229** | ***Ll*ArcA** | **0.256** | ***Ll*ArcB** | **0.087** | ***Ll*ArcC1** | **0.558** | ***Ls*ArcD** |
| **13** | 0.229 | *Ll*ArcA | 0.233 | h*Ll*ArcB | 0.249 | *Ll*ArcC1 | 0.239 | h*Ls*ArcD |
| **18** | 0.429 | mixed | 0.753 | h*Ll*ArcB | 0.599 | *Ll*ArcC1 | 0.594 | h*Ls*ArcD |
| **20** | 0.066 | *Ll*ArcA | 0.727 | h*Pa*ArcB | 0.424 | *Ll*ArcC1 | 0.756 | h*Ls*ArcD |
| **23** | 0.909 | h*Ll*ArcA | 0.073 | h*Pa*ArcB | 0.249 | *Ll*ArcC1 | 0.935 | h*Ls*ArcD |
| **27** | 0.229 | *Ll*ArcA | 0.421 | h*Pa*ArcB | 0.087 | *Ll*ArcC1 | 0.260 | *Ls*ArcD |
| **35** | 0.901 | *Ll*ArcA | 0.401 | h*Ll*ArcB | 0.753 | *Ll*ArcC1 | 0.079 | h*Ls*ArcD |
| **39** | **0.909** | **h*Ll*ArcA** | **0.563** | **h*Pa*ArcB** | **0.599** | ***Ll*ArcC1** | **0.558** | ***Ls*ArcD** |
| **54** | **0.229** | ***Ll*ArcA** | **0.401** | **h*Ll*ArcB** | **0.599** | ***Ll*ArcC1** | **0.239** | **h*Ls*ArcD** |
| **58** | 0.909 | truncated h*Ll*ArcA | 0.909 | - | 0.918 | *Ll*ArcC1 | 0.756 | h*Ls*ArcD |
| **68** | 0.909 | h*Ll*ArcA | 0.107 | *Ll*ArcB | 0.087 | *Ll*ArcC1 | 0.756 | h*Ls*ArcD |
| **73** | - | - | 0.563 | h*Pa*ArcB | mixed | *Ll*ArcC1 | - | Mixed |

| | | | | | | | | |
|---|---|---|---|---|---|---|---|---|
| **76** | 0.901 | *Ll*ArcA | 0.897 | h*Ll*ArcB | 0.918 | *Ll*ArcC1 | 0.935 | h*Ls*ArcD |
| **81** | - | - | 0.075 | h*Ll*ArcB | mixed | *Ll*ArcC1 | - | Mixed |
| **83** | **0.229** | ***Ll*ArcA** | **0.250** | **h*Pa*ArcB** | **0.599** | **h*Ll*ArcC1** | **0.935** | **h*Ls*ArcD** |
| **88** | 0.909 | h*Ll*ArcA | 0.073 | h*Pa*ArcB | 0.599 | *Ll*ArcC1 | - | - |
| **89** | 0.229 | *Ll*ArcA | 0.427 | *Ll*ArcB | 0.753 | *Ll*ArcC1 | 0.594 | h*Ls*ArcD |
| **95** | 0.429 | *Ll*ArcA | - | - | - | - | - | - |
